# Hemispheric laterality of the putamen predicts pseudoneglect

**DOI:** 10.1101/2025.09.17.676561

**Authors:** Tara Ghafari, Mohammad Ebrahim Katebi, Mohammad Hossein Ghafari, Aliza Finch, Kelly Garner, Ole Jensen

## Abstract

Healthy individuals tend to exhibit a subtle leftward attentional bias, a phenomenon termed pseudoneglect. While this bias is thought to reflect a right-hemisphere dominance when allocating spatial attention, the contribution of subcortical structures remains poorly understood. Although lesion and neuroimaging studies in clinical populations have implicated asymmetries in the basal ganglia and thalamus to dysfunction in spatial attention, it remains unclear whether naturally occurring subcortical asymmetries in healthy individuals predict behavioural biases in spatial attention. In this study, we investigated whether volumetric asymmetries in subcortical structures are associated with biases in spatial attention in a non-clinical population. This was achieved by using the landmark task to assess spatial biases, alongside acquiring T1-weighted MRI data from 44 healthy participants. The subcortical regions were segmented using the FIRST algorithm to estimate the volumetric asymmetry of specific regions. We found that variability in pseudoneglect was predicted by the degree of leftward lateralisation of putamen volume, indicating that the putamen contributes to the magnitude of spatial attention biases. This also suggests that the left putamen may support right-hemisphere neocortical dominance through the indirect pathway. These findings bridge anatomical and behavioural measures, highlighting the functional relevance of subcortical asymmetry in shaping attentional processes in the healthy brain. Our findings pave the way for early diagnosis of neurological disorders associated with subcortical atrophies including Parkinson’s Disease and dementia.

**Significance Statement:** Hemispheric dominance in cortical networks shape attentional biases, but the role of subcortical structures remains poorly understood. Here we show that natural variability in the lateralisation of the putamen predicts individual differences in pseudoneglect—a subtle leftward bias in spatial attention observed in healthy adults. Using MRI and psychophysics, we demonstrate that greater leftward putamen asymmetry is associated with stronger leftward bias, independent of eye dominance and microsaccadic behaviour, and moderated by handedness. These findings reveal a structural subcortical basis for attentional asymmetries and suggest that simple behavioural measures could serve as non-invasive markers of early subcortical changes relevant to disorders such as Parkinson’s disease, Alzheimer’s disease, and ADHD.

## Introduction

Healthy individuals typically exhibit a subtle but consistent leftward attentional bias, known as pseudoneglect (Bowers & Heilman, 1980). This phenomenon has been robustly demonstrated using tasks such as manual line bisection (Jewell & McCourt, 2000) and the landmark task (Harvey et al., 1995; Reuter-Lorenz et al., 1990), where healthy participants tend to mark or judge the midpoint of a horizontal line to the left of its true centre. These left hemifield biases are the mirror opposite of the deficits seen in patients with hemispatial neglect caused by right hemisphere lesions (Heilman & Valenstein, 1979). While this spatial attention bias is subtle in healthy adults, it is thought to reflect asymmetrical neural organisation that favours right-hemisphere for attentional control (Bowers & Heilman, 1980; Reuter-Lorenz et al., 1990; Szczepanski & Kastner, 2013).

Although this hemispheric asymmetry has traditionally been attributed to neocortical mechanisms (Çiçek et al., 2009; Corbetta et al., 2005; Corbetta & Shulman, 2011), accumulating evidence highlights the role of subcortical structures—particularly within the basal ganglia and thalamus—in modulating attentional processes (Boshra & Kastner, 2022; Esposito et al., 2021; Saalmann et al., 2012). Lesion studies in patients with spatial neglect have consistently implicated right-sided damage to the putamen, pulvinar, and caudate nucleus to the manifestation of severe attentional deficits (H.-O. Karnath et al., 2004). Neuroimaging work in neurodivergent populations, such as individuals diagnosed with ADHD, has revealed atypical asymmetries in subcortical structures including the globus pallidus, caudate, and putamen, as compared with healthy controls (Hynd et al., 1993; Uhlíkova et al., 2007; Wellington et al., 2006; but also see Postema et al., 2021). Moreover, in neurodegenerative disorders such as Alzheimer’s disease, the asymmetry of subcortical regions is altered, which is correlated with measures of cognitive assessment tests (Fu et al., 2021). Importantly, even among healthy adults, subclinical attention impairments associated with ADHD symptomatology have been correlated with caudate nucleus asymmetry (Dang et al., 2016).

In non-clinical populations the functional significance of subtle volumetric asymmetries in subcortical volumes remains underexplored. Large-scale morphometric analyses have revealed systematic hemispheric differences in subcortical volumes across thousands of healthy individuals, influenced by factors such as age, sex, and genetics (Guadalupe et al., 2016). Yet, it remains unknown how these anatomical findings link to behavioural measures of spatial attention. This represents a critical gap in our understanding of how structural brain asymmetries relate to attentional function.

Recent findings suggest that individual differences in subcortical structure may influence neuronal activity associated with spatial attention. In our previous work we showed that lateralisation of caudate and globus pallidus volumes predicted hemispheric asymmetry in the modulation of alpha oscillations during an attention task (Ghafari et al., 2024). Similarly, Mazzetti et al. (2019) applied the same analytical approach in a reward-based attention task and found that asymmetries in the globus pallidus and thalamus were linked to lateralized alpha power. Notably, the association between globus pallidus asymmetry and alpha modulation was particularly evident in trials where visual stimuli carried high-value reward valence, whether positive or negative. These findings imply that structural asymmetries in the basal ganglia could underlie lateralised neural dynamics associated with attention. While the association between lateralization in subcortical regions and the modulation of alpha oscillations is now established, it remains unknown if these subcortical asymmetries also correspond to behavioural biases in spatial attention.

The present study investigates whether naturally occurring asymmetries in subcortical structures are associated with spatial attention biases in healthy adults, as occurs in pseudoneglect. Considering anatomical and functional evidence, we hypothesise that asymmetries in the thalamus and basal ganglia structures contribute to individual differences in pseudoneglect. We used the landmark task, a validated measure of spatial attention bias, to quantify pseudoneglect (Strappini et al., 2023; Szczepanski & Kastner, 2013). Using T1-weighted MRI we quantified lateralised volumes of subcortical structures including the putamen, caudate, globus pallidus, nucleus accumbens, hippocampus, amygdala and thalamus. This approach allows us to test whether volumetric asymmetries in these regions are functionally related to the direction and magnitude of spatial bias.

## Method and Materials

### Participants

The research complied with the ethical standards of the Declaration of Helsinki and was approved by the University of Birmingham’s STEM Ethical Review Committee (ERN_18-0226AP37). Participants were recruited from a pool of individuals who had previously undergone a 3T MRI scan as part of other studies. Forty-five volunteers participated; one was excluded due to poor behavioural performance (see Behavioural analysis below). The remaining 44 participants (age range: 21–36 years; mean age: 25.1 years; 31 females; 1 left-handed) had no diagnosed neurological or psychiatric disorders and had normal or corrected-to-normal vision. All participants provided written informed consent prior to the study and were compensated at a rate of £7 per hour.

To consider the potential influence of hand and eye dominance on spatial attention biases (Roth et al., 2002), all participants were tested for both. Handedness was assessed using the Edinburgh Handedness Inventory (Oldfield, 1971), in the version adapted by M. Cohen (Staglin IMHRO Center for Cognitive Neuroscience, UCLA; http://www.brainmapping.org/shared/Edinburgh.php). Eye dominance was identified using a modified Porta test (originally described by Porta (Della Porta, 1593)).

### Landmark Task and procedure

To assess behavioural spatial bias, participants completed a computerized version of the standard *line bisection* (or *landmark*) task, a well-established measure in neuropsychological assessment (Szczepanski & Kastner, 2013). In each trial the stimuli were composed of a horizontal white line bisected by a short vertical line (2° of visual angle) on a grey background. The bisection mark was aligned with the participant’s head and body midline, with the horizontal line displayed at eye level. All lines were 0.1° thick, and the head position was stabilized using a chinrest to ensure consistent viewing. The experimental paradigm was programmed and executed on a Windows 10 system using MATLAB (MathWorks Inc., Natick, MA, USA) in combination with Psychophysics Toolbox version 3.0.11(Brainard, 1997; Pelli, 1997).

Stimuli varied across four different horizontal line lengths (ranging from 20° to 23° of visual angle), presented in random order. Line length variation was included to minimize biases associated with fixed-length stimuli, as longer lines have been shown to affect bisection performance (Anderson, 1997; Bisiach et al., 1983). For each trial, a fixation cross appeared for a random duration between 1000 to 2000 ms, followed by the transected line for 200 ms—a duration brief enough to discourage saccadic eye movements. This was immediately followed by a 2000 ms full-screen pixelated mask, during which participants indicated their response.

In a block design, participants were instructed to perform one of two tasks: to judge which side of the horizontal line was longer, or which side was shorter, depending on the block instruction. Responses were made using the right index, middle, or ring finger to indicate “left,” “neutral” (i.e., both sides perceived as equal), or “right,” respectively. The order of task instructions was counterbalanced across subjects (Figure 1).

**Figure 1.**
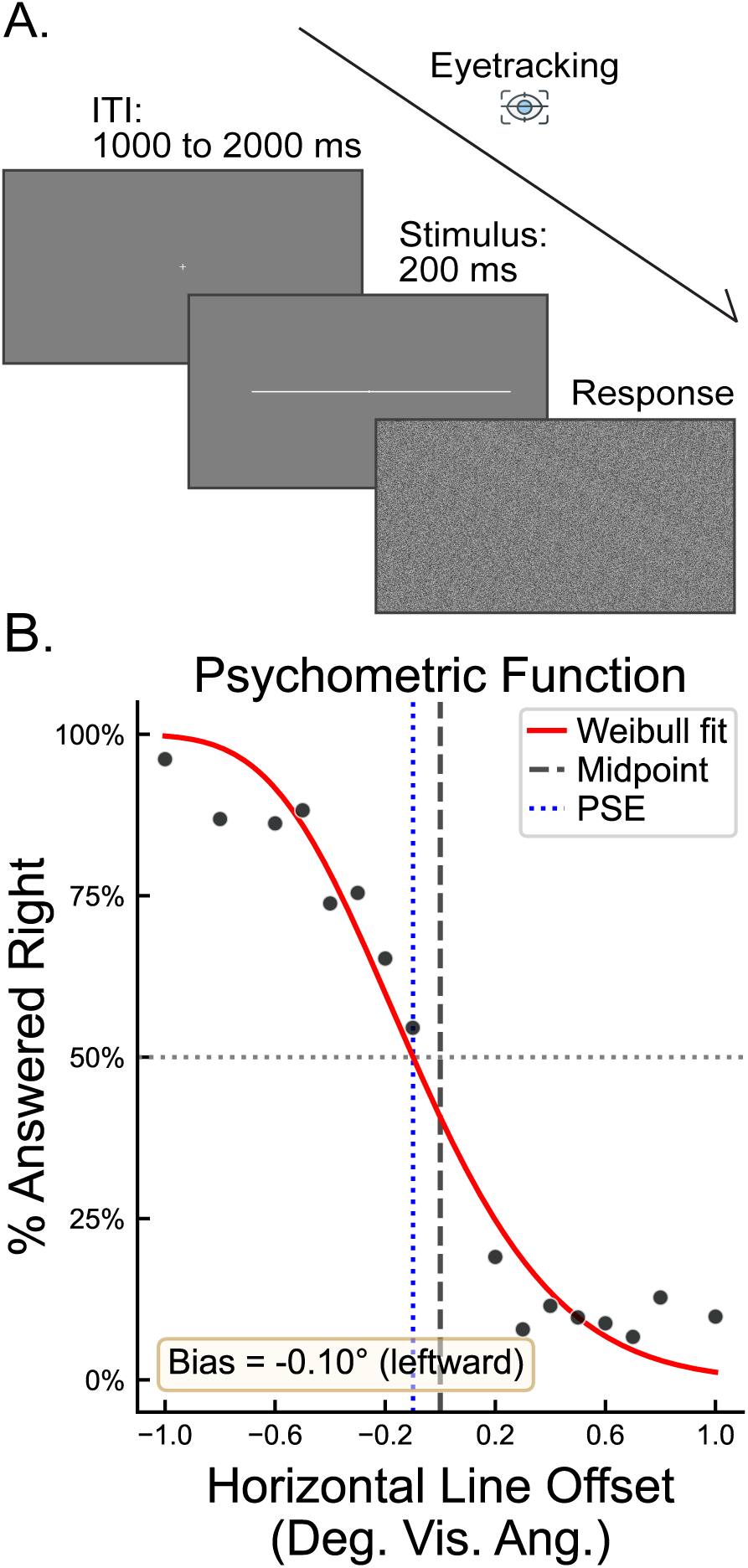
The Landmark Task. A) Participants were instructed to judge which side of a horizontal line appeared shorter or longer, depending on the block’s instruction. Each trial began with a fixation point (1000–2000 ms), followed by a horizontal line with a central vertical transector, presented for 200 ms. The horizontal line was offset between 1° left and 1° right of centre. B) Psychometric function from a representative participant, with a point of subjective equality (PSE) of −0.1, indicating a leftward bias. PSE = Point of Subjective Equality; Deg. Vis. Ang. = Degree of Visual Angle

A staircase procedure was used to determine the point of subjective equality (PSE); i.e. the offset at which participants perceived the two sides as equal in length. Each block began with lines offset by 1° of visual angle either to the left or right of fixation. The staircase followed a two-down, one-up rule: after two consecutive correct responses at a given offset, the line was shifted closer to the midline, with the new offset set to 80% of the previous one. After one incorrect or “neutral” response, the offset was increased to the previous step (i.e., shifted away from the midline), making the task easier. Staircases were run from both the left and right sides, with trials randomly intermixed.

Each block consisted of 240 trials, with four blocks per session: two blocks requiring “longer” judgments and two requiring “shorter” judgments. The entire task lasted approximately 50 minutes, with three brief breaks between blocks to reduce fatigue.

### Behavioural analysis

Trials from each block were organized into bins on a linear scale based on their distance from the veridical midpoint, with bin boundaries set at 0.1° of visual angle apart. For each bin within a single block, we calculated the percentage of times participants responded “right is longer.” For blocks with instruction “Which side is shorter?”, the answers were reversed so that “left is shorter” counted as “right is longer.” These data points were then plotted against the centre offset of each bin where the x-axis showed the offset to the left or right, and the y-axis showed the percent of “right is longer” responses. Thus, each point on the graph represents one bin from one of the four blocks. Finally, to estimate each participant’s spatial bias, a Weibull function was fitted to the data using the SciPy library in Python. The point of subjective equality (PSE) —where left and right responses occurred with equal frequency— was then derived from the psychometric function for each individual.

Note that PSE measures have been reported in different ways in the literature. In the classical line bisection task, a negative PSE indicates a mark to the left of the centre of the line (Bradshaw et al., 1987; Harvey et al., 1995). This is also interpreted as a left hemifield bias. An alternative approach calculates the PSE based on the horizontal offset required (left or right) for the line’s centre to be perceived as midpoint (Szczepanski & Kastner, 2013); under this convention, a left hemifield bias is reflect by a positive PSE. While we follow the method of Szczepanski & Kastner, 2013, we report PSE values according to the classical line bisection task (i.e., a negative PSE reflects a shorter bisection to the left than to the right).

We then plotted the distribution of behavioural biases among the participants (figure 2). This depicted the number of participants with right (RVF) or left (LVF) biases relative to the veridical midpoint to determine behavioural lateralisation.

**Figure 2.**
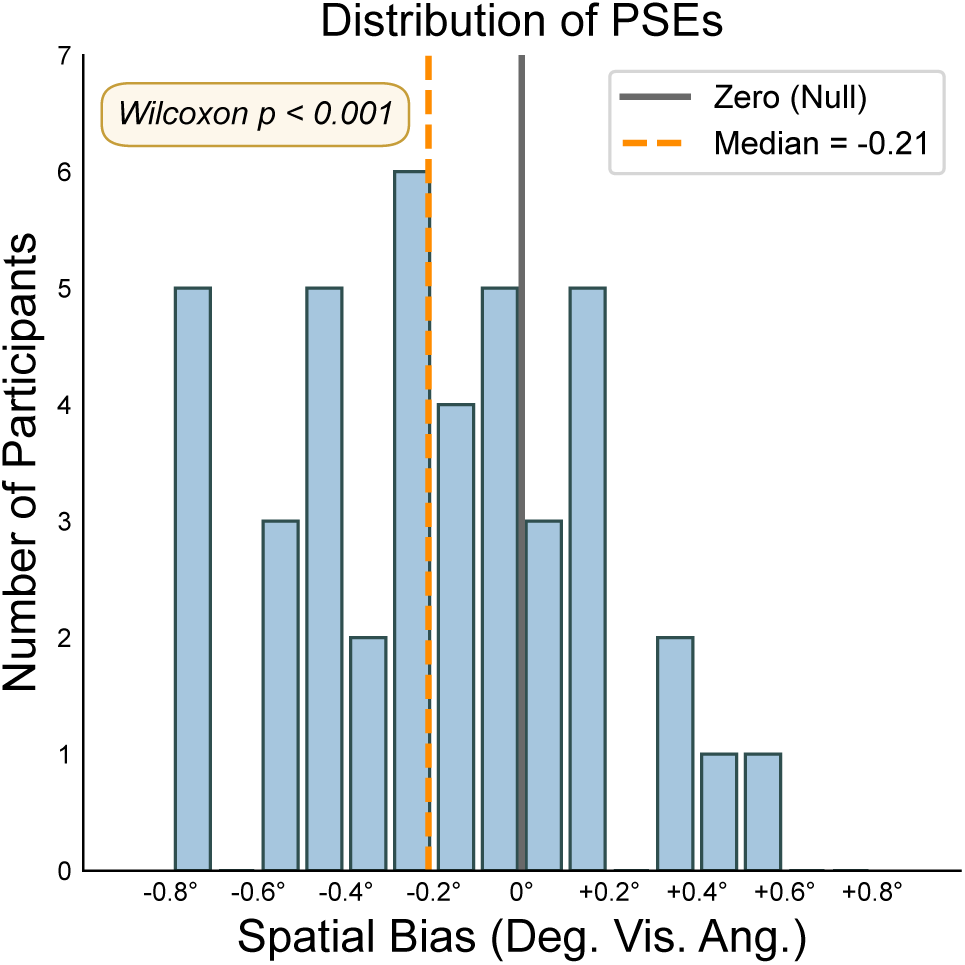
Distribution of point of subjective equality (PSE) values from the landmark task across all participants. The histogram depicts the frequency of PSE values indicating spatial biases, with positive values reflecting a rightward bias and negative values indicating a leftward bias. The vertical lines represent zero (green), and median (orange). A Wilcoxon rank sum test showed that the group median was significantly different from zero (p = 0.0004), indicating a significant leftward bias in spatial perception across the sample (median = −0.21), whereby participants judged lines with a physically longer right side as appearing symmetrical. PSE = Point of Subjective Equality; Deg. Vis. Ang. = Degree of Visual Angle

### Eye Tracking Data Analysis

#### Data Acquisition

Eye movements were recorded binocularly using an EyeLink eye tracker (EyeLink 1000, SR Research Ltd., Ottawa, Canada) with a sampling rate of 500 Hz. The eye tracker was positioned approximately 5 cm in front of the monitor and 60 cm away from the participant’s eyes. Prior to recording, the built-in calibration and validation protocols from the EyeLink software were used to ensure accurate gaze tracking. The raw data were converted to ASCII format using EyeLink’s *edf2asc* utility. All subsequent analyses were performed using custom Python scripts.

#### Data Preprocessing and Segmentation

The eye tracking data were segmented into epochs corresponding to individual trials. Each epoch began at stimulus onset and extended for 1700 ms, encompassing the 200 ms stimulus duration and 1500 ms post-stimulus offset. Blinks were identified using a velocity-based algorithm adapted from Hershman et al. (2018). To account for potential artifacts, 40 ms of data before and after each blink were also excluded.

#### Data Filtering and Quality Control

Trials were excluded from the analysis if the participant’s gaze deviated more than 2° from the central fixation point for more than 10% of the trial duration. This criterion ensured that only trials with stable fixation were included in the analysis.

#### Microsaccade Detection and Characterization

Microsaccades were detected using a velocity-based algorithm adapted from Engbert and Kliegl (2003). Gaze velocity was derived by computing the absolute temporal derivative of the gaze position. To reduce noise, the resulting velocity trace was smoothed using a Gaussian-weighted moving average filter with a 7-ms window.

The algorithm calculated velocity thresholds for each trial separately to account for trial-specific noise levels. Saccades were identified as periods where the velocity exceeded a threshold of 5 times the median-based standard deviation of the velocities. Only saccades observed binocularly (i.e., saccades detected in both eyes with at least 2 overlapping data points) were considered for further analysis.

To prevent multiple counts of a single eye movement, a minimum interval of 100 ms was required between consecutive gaze-shift initiations. Only saccades with amplitudes less than 1° were classified as microsaccades. We further categorized microsaccades as either “start microsaccades” (moving away from fixation) or “return microsaccades” (moving back toward fixation) by comparing pre- and post-saccade distances from the fixation position.

#### Microsaccade Laterality Index

To quantify the overall directional bias of microsaccades for each participant, we defined a Microsaccade Laterality Index (MLI). The MLI was computed as (R - L) / (R + L), where R is the number of rightward microsaccades (315° to 45°), and L is the number of leftward microsaccades (135° to 225°). This index ranges from −1 (strong leftward bias) to +1 (strong rightward bias). To prevent the return microsaccades (i.e., saccades with trajectories towards fixation) cancelling out the start microsaccades (i.e., saccades with trajectories away from fixation) (Liu et al., 2023), we only included start microsaccades in our analysis. This approach yields a more accurate overall measure of microsaccade behaviour relative to the central fixation point.

### MRI data acquisition and analysis

Structural MRI data (T1-weighted) were sourced from prior studies in which all participants had undergone scanning using a 3 Tesla Siemens Magnetom Prisma whole-body scanner. The imaging parameters were as follows: repetition time (TR) = 2000 ms, echo time (TE) = 2.01 ms, inversion time (TI) = 880 ms, field of view (FoV) = 256 × 256 × 208 mm³, and an isotropic voxel size of 1 × 1 × 1 mm.

Subcortical segmentation was carried out using FMRIB’s Integrated Registration and Segmentation Tool (FIRST), version 5.0.9, available through the FSL software package (https://www.fmrib.ox.ac.uk/fsl/). This tool applies a model-based approach within a Bayesian framework, incorporating shape and intensity priors derived from manually labelled datasets (Centre for Morphometric Analysis, MGH, Boston). An initial affine registration (FLIRT) aligns each participant’s scan to MNI152 standard space, after which FIRST identifies subcortical structures including the thalamus, caudate, putamen, globus pallidus, hippocampus, amygdala, and nucleus accumbens. Each structure is represented as a surface mesh fitted to the subject’s original space, with non-subcortical voxels excluded from the segmentation process (Patenaude et al., 2011).

To quantify asymmetry between hemispheres, we computed lateralization volume indices (LVs) for each structure:

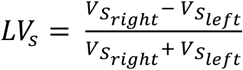

where 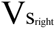 and 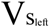 denote the voxel-based volume of a subcortical regions in the right and left hemispheres, respectively. This metric accounts for individual variability in overall brain size and is commonly employed in studies of structural asymmetry (e.g., Ghafari et al., 2024). LV values range from –1 to 1, with positive values reflecting rightward asymmetry and negative values indicating leftward asymmetry.

### Statistical analysis

#### Model selection

To examine how individual differences in spatial perception bias, indexed by the point of subjective equality (PSE), relate to asymmetries in subcortical brain structures, we conducted a series of generalized linear model (GLM) analyses.

PSE values served as the dependent variable, while LVs of subcortical regions were included as explanatory variables. Prior to conducting the GLM, we assessed multicollinearity among predictors using the *statsmodels* library to ensure sufficient independence between them. No predictor had a VIF values above 5, so we included all of them in the GLM model.

Our primary goal was to determine the combination of predictors that most effectively accounted for variance in PSE. To this end, we systematically tested all possible subsets of the seven subcortical lateralization indices (LVs), including interaction terms between basal ganglia structures, in order to assess whether combined anatomical asymmetries exerted synergistic effects on spatial bias. Model comparisons were conducted using ordinary least squares regression, spanning from single-variable to multivariate combinations. The optimal model was selected based on the lowest Akaike Information Criterion (AIC; Akaike, 1974), with the Bayesian Information Criterion (BIC; Schwarz, 1978) used as a confirmatory metric. Both criteria supported similar model selections, balancing model fit against complexity. In our analysis, the winning model included the lateralized volume of the putamen (LV_put_) as the sole predictor. The final model was as follows:

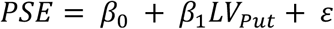

where PSE is the behavioural index of spatial bias and LV_Put_ refers to the lateralised volume of the putamen, *β*_0_ and *β*_1_ are the model intercept and slope, respectively, and *ɛ* is the residual error term.

#### Cross-Validation

Next, we sought to determine the accuracy with which the best model would predict new samples. Specifically, we used two convergent measures that each estimate the R^2^ value that would be obtained if the model was used to predict new samples (cross-validation correlation coefficient). First, we analytically calculated the cross-validity correlation coefficient using the following formula (Pedhazur, 1997):

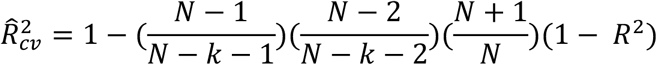

where *N* is the number of participants, and *k* is the number of predictors (in our case 1 for LV_put_) and *R*^2^ is the observed squared multiple correlation. Using an analytical means to estimate the cross-validation correlation coefficient supports interpretation of model generalisability when there exist limited means to perform a cross-validation analysis, such as when the model has been estimated using a limited sample (Cotter & Raju, 1982).

Next, we used the data to estimate the cross-validity correlation coefficient, by applying a split-half cross-validation. The dataset was randomly divided into two parts: one subset (the screening sample) was used to estimate the regression equation, and this equation was then applied to the other subset (the calibration sample) to generate predicted values. We then compared these predicted values with the actual observed scores in the calibration sample using a Pearson correlation.

#### Mediation and Moderation Analyses

We conducted mediation and moderation analyses to explore whether variability in handedness, eye dominance, or microsaccade laterality (control variables) could explain or influence the winning model. Mediation analysis followed the classic three-step procedure outlined by Baron and Kenny (1986):

- Total effect: We regressed PSE on the winning LV predictor (LV_Put_) to establish a total effect:

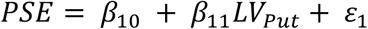
- Path a: We regressed control variable (*Var*_*C*_) on LV_Put_ to test whether the independent variable predicted the mediator:

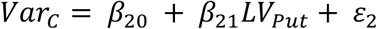
- Path b and c’: We then regressed PSE on both the LV_Put_ and MLI to assess the mediator’s contribution to the relationship.

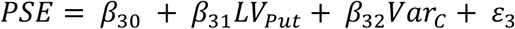

The indirect effect was calculated as the product of the coefficients from paths a and b (*β*_21_ ∗ *β*_32_). Significance was evaluated using bootstrapped confidence intervals.

Finally, to test for moderation, we included interaction terms between the LV_Put_ and *Var*_*C*_. This assessed whether the relationship between anatomical asymmetry and spatial bias varied as a function of the control variable.

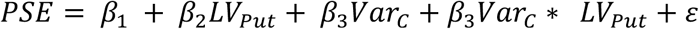

Mediation and Moderation analysis were run separately on the three control variables. All statistical procedures were performed using Python libraries including *pingouin*, *seaborn*, and *statsmodels*. Visualisation of variable relationships and diagnostics was aided by Matplotlib, *pairplots* and residual plots to ensure model validity.

### Data and code availability

The code used for the experiment, along with the data and analysis scripts, can be found at:https://github.com/tghafari/behavioral_structural_asymmetry”

## Results

### A significant leftward bias in line bisection judgments

To assess participants’ perceptual spatial bias, we analysed the distribution of point of subjective equality (PSE) values derived from the landmark task. The histogram in Figure 2, shows the number of participants in relation to individual PSE values, with positive values indicating a right hemifield bias and negative values indicating a leftward bias. The group-level distribution exhibited a significant leftward bias, as reflected by a median PSE of −0.21. A Wilcoxon rank sum test confirmed that the median PSE was significantly different from zero (p = 0.0004), indicating a robust overall bias toward the left visual hemifield. This pattern suggests overrepresentation or enhanced perceptual weighting of the left side of space, requiring the right segment of the line to be physically longer in order to perceive both sides as equal.

### Subcortical brain structures show consistent hemispheric asymmetries

To investigate structural asymmetries in subcortical regions, we calculated lateralisation volume (LV) indices for each participant using T1-weighted MRI scans. As illustrated in Figure 3 and in line with our previous work (Ghafari et al., 2024), the group-level distribution of LVs revealed consistent hemispheric asymmetries across several subcortical structures. Specifically, the thalamus (median LV = −0.0104, Wilcoxon rank sum p-value < 0.001), putamen (median LV = −0.0132, Wilcoxon rank sum p-value < 0.001), and nucleus accumbens (median LV = −0.0847, Wilcoxon rank sum p-value < 0.001) exhibited a leftward lateralization pattern, indicating larger volumes in the left hemisphere. In contrast, the caudate nucleus (median LV = 0.0163, Wilcoxon rank sum p-value < 0.001) showed a rightward bias, with greater volume in the right hemisphere. These patterns suggest a systematic anatomical lateralization of subcortical structures at the population level. Globus Pallidus (median = −0.0009, Wilcoxon rank sum p-value = 0.65), Hippocampus (median = 0.0098, Wilcoxon rank sum p-value = 0.34), and Amygdala (median = 0.0053, Wilcoxon rank sum p-value = 0.49) did not show asymmetry in our sample.

**Figure 3.**
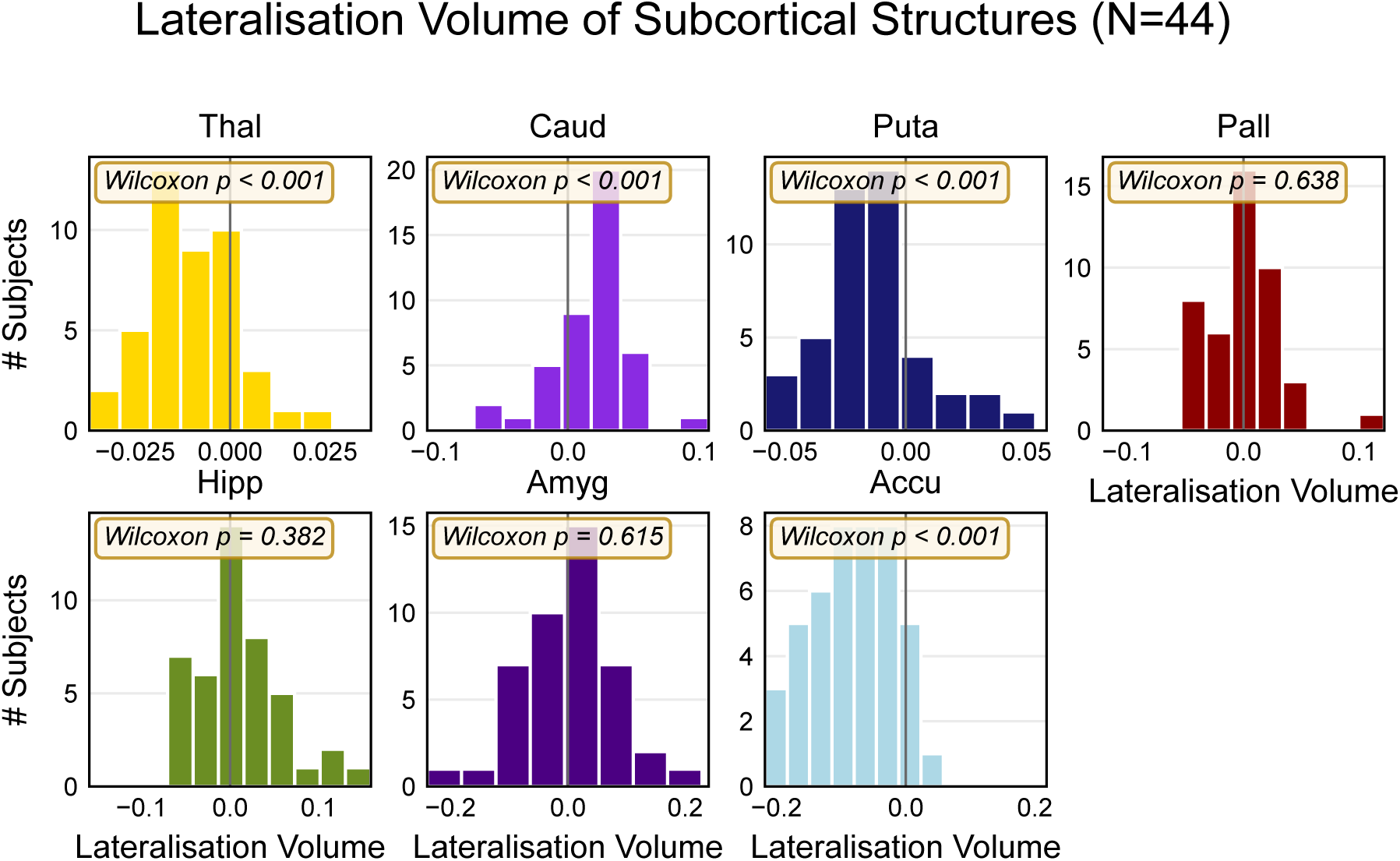
Distribution of lateralisation volume indices for subcortical structures across participants. Positive values indicate greater volume in the right than left hemisphere and vice versa for negative values. The thalamus, putamen, and nucleus accumbens show consistent leftward asymmetry, whereas the caudate nucleus demonstrates rightward asymmetry. Thal = Thalamus; Caud = Caudate nucleus; Puta = Putamen; Pall = Globus pallidus; Hipp = Hippocampus; Ayg = Amygdala; Accu = Nucleus accumbens

### Greater left-lateralisation of the putamen predicts increased leftward perceptual bias

To determine whether volumetric asymmetry of subcortical structures could account for variability in spatial perceptual bias, we conducted a linear regression analysis using the lateralisation indices of subcortical volumes as predictors of landmark task PSE values. Model comparison revealed that the best-fitting model included only the lateralisation index of the putamen (AIC = 33.6, BIC = 37.2, formula-based cross-validity coefficient = 0.070, half-split cross-validity coefficient = 0.32) (See Supplementary Materials for a full table of AICs and BICs).

The regression coefficient for the putamen was positive (β = 6.2, SE = 2.4), with a 95% confidence interval of [1.26, 11.1] (Figure 4B). This result indicates that participants with greater leftward volumetric asymmetry in the putamen (i.e., larger left than right hemisphere putamen) exhibited a stronger leftward bias in spatial perception, as indexed by more negative PSE values. This relationship suggests a functional link between the anatomical asymmetry of the putamen and hemifield biases in visual perception.

**Figure 4.**
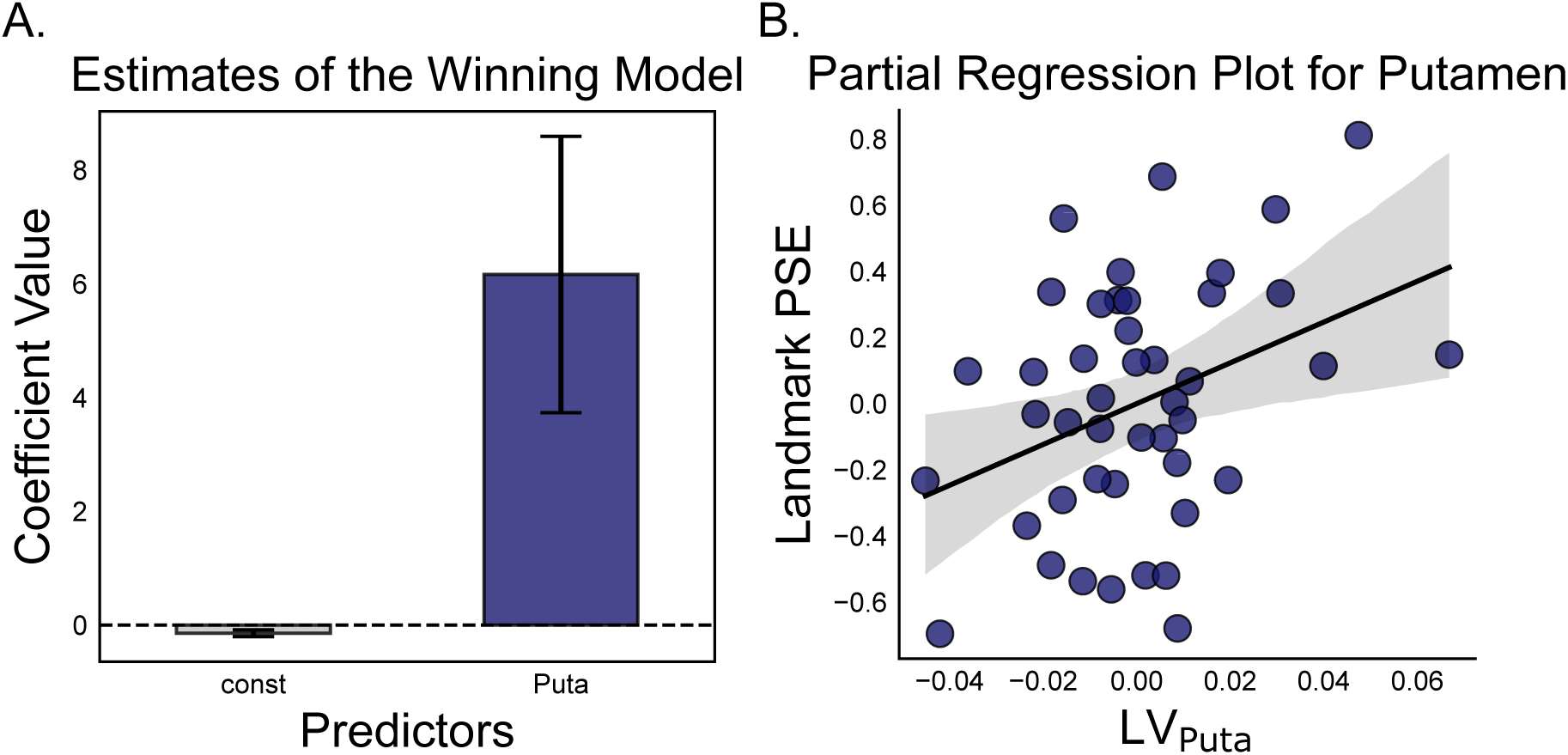
A) Bar plot showing the beta coefficient of the model explaining the relationship between putamen lateralisation index and landmark PSE values. The regression model ((β = 6.2, SE = 2.4, 95% confidence interval = [1.26, 11.1]) indicates that greater leftward asymmetry of the putamen is associated with a more pronounced leftward perceptual bias. B) Partial regression plot, isolating the effect of putamen lateralisation on the dependent variable. The shaded area represents the 95% confidence interval of the regression line. PSE = Point of Subjective Equality; LV = Lateralised Volume; Puta = Putamen

### Handedness Moderates—but Does Not Mediate—the Relationship Between Putamen Asymmetry and Spatial Attention Bias; Microsaccade Laterality and Eye Dominance Show No Mediation or Moderation Effects

To determine whether individual differences in handedness, eye dominance, or microsaccade laterality might account for the observed relationship between putamen asymmetry and spatial attention bias, we conducted mediation and moderation analyses for each variable. Mediation analysis tested whether the control variable provided an explanatory link between putamen asymmetry and participants’ PSE scores, while moderation analysis assessed whether this relationship varied depending on hemifield biases in microsaccades, dominant eye or handedness. Microsaccade laterality and eye dominance showed no significant effects in either analysis (microsaccade laterality mediation: *p* = 0.494; moderation: Adjusted R² = 0.070, model *p* = 0.127; eye dominance mediation: *p* = 0.920; moderation: Adjusted R² = 0.029, model *p* = 0.262), suggesting they do not contribute to or modify this relationship (Supplementary figure 1 and table 2). In contrast, handedness showed a significant moderation effect (Adjusted R² = 0.260, model *p* = 0.002): the model containing handedness, putamen and the interaction between the two is significantly correlated with landmark PSE scores, indicating that the relationship between putamen asymmetry and behavioural bias is stronger in individuals with larger handedness scores (strongly right-handed individuals). In this model, the regression coefficient for putamen asymmetry was significantly positive (β = 35.0, SE = 12.32, CI = [0.29, 1.45], *p* = 0.007), while the coefficient for the main effect of handedness (β = −0.01, SE = 0.00, CI = [−0.02, −0.00], *p* = 0.004) and the putamen– handedness interaction were significantly negative (β = −0.34, SE = 0.14, CI = [−0.62, −0.05], *p* = 0.025) (Figure 5A). However, no mediation effect (mediation: *p* = 0.196) was observed for handedness, suggesting that even though handedness modulates, it does not explain the association between putamen asymmetry and attentional bias.

**Figure 5.**
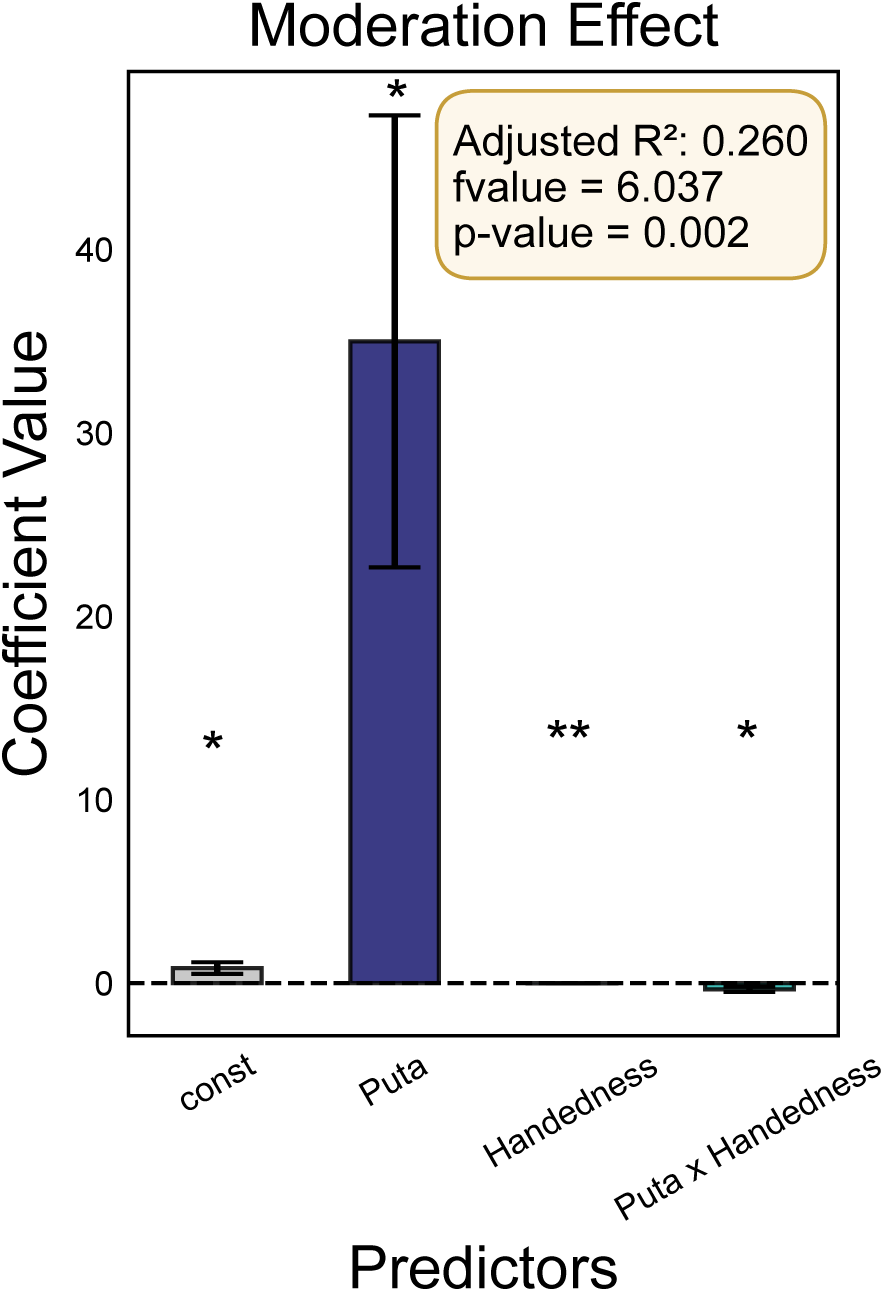
Moderation effect of handedness on the relationship between putamen asymmetry and landmark PSE. The regression model which included handedness, putamen asymmetry, and their interaction, showed a significant association with landmark PSE scores, suggesting that the strength of the putamen-attention bias relationship varies as a function of handedness. Adjusted R² = 0.260, model *p* = 0.002. Puta = Putamen; single asterisk (*) denotes p < 0.05; double asterisks (**) denote p < 0.005.

## Discussion

Using the landmark task in 44 participants, we replicated a bias to the left hemifield: when judging both sides of a line equal, the right element was in fact longer than the left. The magnitude of this leftward bias varied systematically with basal ganglia anatomy: individuals with greater leftward volumetric asymmetry of the putamen (i.e. a larger left than right putamen) showed a stronger leftward perceptual bias in the landmark task. Control analyses revealed that handedness influences the strength of the relationship between putamen asymmetry and pseudoneglect, acting as a moderator but not a mediator. Neither microsaccade lateralisation nor eye dominance showed any moderating or mediating effects on this association.

Our findings provide new evidence linking perceptual biases in terms of hemifield lateralisation to structural asymmetries in subcortical structures. The observed leftward bias in the landmark task, mimicking the land-bisection task, aligns with pseudoneglect, where healthy participants show a bias toward the left visual field by judging a longer right side of the line to achieve perceptual balance (Bowers & Heilman, 1980; Jewell & McCourt, 2000; Reuter-Lorenz et al., 1990). This tendency is commonly attributed to a right-hemispheric dominance in spatial attention networks (Heilman & Abell, 1980; Mesulam, 1981). Critically, the association between greater leftward asymmetry of the putamen and enhanced leftward bias suggests that the basal ganglia—specifically the putamen— biases the fronto-parietal attention work possibly via connections to neocortex mediated by the anterior thalamus (Alexander et al., 1986; Di Martino et al., 2008; Haber, 2003; Jarbo & Verstynen, 2015). Evidence from healthy individuals supports this view: eye-blink rate, a proxy for striatal dopamine (Slagter et al., 2010), asymmetries in dopaminergic neurotransmission (Tomer et al., 2013), and genetic variation affecting dopamine function (Zozulinsky et al., 2014) all predict the magnitude of pseudoneglect and orienting biases. These findings highlight dopamine lateralisation as a potential neurochemical mechanism linking structural asymmetry of the putamen to attentional biases. We suggest that this effect is more likely mediated by the striatal indirect rather than the direct pathway, given that the indirect pathway exerts inhibitory control over neocortex. Supporting this, an optogenetic study in mice found that activating the striatal direct pathway (which includes the putamen in humans) shifted decision bias in a visual attention task, specifically, when attention was directed to the contralateral side of the stimulated striatum (Wang & Krauzlis, 2020). While these findings show that the basal ganglia can modulate perceptual decisions in a hemifield specific manner, further work is required to uncover the specific pathways of action in relation to the direct and indirect pathways.

We found that handedness significantly moderated—but did not mediate—the relationship between putamen asymmetry and spatial attention bias. This finding aligns with prior research showing that right-handed individuals tend to bisect lines further to the left than left-handed individuals (Luh, 1995; Scarisbrick et al., 1987). Handedness is known to reflect individual differences in hemispheric lateralisation, which can influence functional brain organisation even in the absence of structural differences (Tomasi & Volkow, 2024). While the moderation effect suggests that handedness influences the strength of the putamen– attention bias relationship, the lack of mediation indicates that this relationship is not driven by handedness itself.

Our findings link structural asymmetries of the putamen in healthy adults to subtle biases in spatial attention, extending prior evidence from clinical and neuroimaging studies. Notably, subcortical structures like the putamen are part of the networks interacting with neocortex whose disruption can produce hemispatial neglect. Strokes affecting the right putamen, pulvinar, and caudate nucleus have been shown to induce neglect possibly by dysregulating a distributed attention network including the superior temporal, inferior parietal, and frontal cortices (Karnath et al., 2002; Karnath et al., 2004). This supports the view that the basal ganglia contribute to spatial awareness alongside cortical structures. Reverse asymmetry of the putamen, where the left putamen is smaller than the right, have been implicated in ADHD and is thought to reflect dysfunctions within the fronto-striatal attention network. Individuals with ADHD typically show reduced pseudoneglect or even a rightward attentional bias (Wellington et al., 2006). Our results extend these observations, suggesting that even in non-clinical populations, individual differences in putamen, asymmetry relates to spatial attention biases. However, large-scale studies such as those conducted by Postema et al. (2021) have sometimes failed to find consistent subcortical asymmetry–behaviour relationships in neurodevelopmental disorders, underscoring the heterogeneity of structural findings. As such, it would be of great interest to conduct a study in a population diagnosed with ADHD to investigate whether difference in spatial biases as measured by the landmark task is linked to structural changes in the putamen. Our findings are consistent with evidence of visual field asymmetries, such as enhanced recognition for stimuli in the upper and right quadrants (Ghafari et al., 2023), showing that perceptual biases extend beyond line bisection tasks.

Further evidence for the role of the putamen in attentional control comes from neurodegenerative diseases. In Parkinson’s disease, early dopaminergic depletion in the posterior putamen disrupts basal ganglia circuits that support automatic visual selection, contributing to subtle visuospatial attention deficits even in early disease stages (Fornari et al., 2021). Similarly, although Alzheimer’s disease is most strongly associated with hippocampal and cortical atrophy, reduced volumes of the putamen and thalamus have also been observed (de Jong et al., 2008).

Crucially, subcortical volume reductions have been shown to correlate with poorer cognitive performance, suggesting that degeneration in these regions contributes to the broader cognitive phenotype observed in both Parkinson’s and Alzheimer’s disease. In Alzheimer’s, attentional deficits often emerge early and may worsen memory encoding processes (Malhotra, 2019), raising the possibility that structural changes in regions such as the putamen play a role in these early impairments. Moreover, as many major neurological disorders exhibit early pathological changes in subcortical regions (Mak et al., 2014; Roh et al., 2011; Sabuncu et al., 2011), the early detection of subcortical abnormalities becomes crucial, offering a window for intervention when therapeutic strategies are likely to be most effective. Recent neuroimaging studies further suggest that subcortical degeneration in both Alzheimer’s and Parkinson’s disease often occurs asymmetrically (Djaldetti et al., 2006; Jahanshahi et al., 2023; Yang et al., 2023), and that lateralised atrophy or shape deformation is associated with specific clinical outcomes, including cognitive and motor deficits (Fiorenzato et al., 2021). These findings imply that subcortical asymmetry is not simply a by-product of global atrophy but may have functional significance in the manifestation and progression of neurodegenerative syndromes.

Several limitations of this study should be considered. Given that our findings are based on correlations, the link between hemifield biases and the putamen might not be causal. It is also possible that long-term biases in sensory experience could shape striatal morphology. The causal role of the putamen in spatial attention could be tested by using brain stimulation techniques such as transcranial focused ultrasound stimulation which can target subcortical regions (Folloni et al., 2019). Additionally, the use of volumetric MRI provides only a gross structural measure and does not reveal the functional or neurochemical status of these subcortical circuits. For example, our approach cannot identify the specific neural populations or neurotransmitter systems within the striatum (e.g. dopaminergic pathways) responsible for the observed effects, nor can it determine whether these effects are primarily mediated by the direct or indirect pathway. These questions could be explored with functional neuroimaging studies in healthy participants or subcortical recordings and optogenetic manipulations in animal models.

Furthermore, the cross-validated coefficients from the winning model ranged from 0.07 to 0.32. This variance in estimates may be due to the current sample size. Nonetheless, even at the lower bound, the model accounted for a meaningful proportion of variability. It should also be noted that, although we controlled for handedness and eye-dominance, other factors governed by hemispheric laterality (e.g. language dominance, subclinical sensorimotor asymmetries) were not assessed.

Future research could extend our findings by combining multiple converging approaches. Behaviourally, it will be informative to test whether individual differences in putamen asymmetry bias performance on other spatial tasks, such as visual search or orienting tasks, and whether similar asymmetry–bias relationships exist in other sensory modalities. Neuroimaging studies using functional MRI or diffusion tensor imaging (DTI) could examine whether structural or functional asymmetries in basal ganglia–thalamo–cortical connectivity align with volumetric asymmetries we observed. Longitudinal designs can test whether these subcortical asymmetries change across development or with training. Causally, non-invasive brain stimulation (such as transcranial focused ultrasound or deep brain stimulation) could be used to transiently alter activity in the basal ganglia, testing for induced changes in spatial bias. Likewise, pharmacological manipulation of neurotransmitter systems (e.g., dopaminergic drugs) could shed light on the modulatory role of these regions on perception and spatial attention.

Finally, studying clinical groups with known basal ganglia dysfunction – for example, patients with unilateral striatal strokes, individuals with Parkinson’s or Huntington’s disease, or even in neurodevelopmental disorders like ADHD with our paradigm could reveal whether the same putamen–bias relationship operates when networks are altered. By integrating structural, functional, and interventional methods, future work can more fully elucidate the role of basal ganglia–thalamo-cortical loops in shaping human spatial attention.

In conclusion, our findings demonstrate that individual differences in putamen hemispheric asymmetry are systematically related to subtle biases in spatial attention. By demonstrating that a simple perceptual bias correlates with putamen asymmetry, our study highlights a potential non-invasive method for identifying emerging structural abnormalities in subcortical areas. In turn, this supports a broader framework in which subcortical asymmetries may serve as early indicators of a wide spectrum of neurological conditions— from dementias such as Alzheimer’s disease to movement disorders including Parkinson’s and Huntington’s disease—many of which begin with subcortical involvement.

## Acknowledgments

This study was funded by Wellcome Trust Discovery Award (grant number 227420) and supported by the NIHR Oxford Health Biomedical Research Centre (NIHR203316). The views expressed are those of the author(s) and not necessarily those of the NIHR or the Department of Health and Social Care. The Oxford University Centre for Integrative Neuroimaging was supported by core funding from the Wellcome Trust (203139/Z/16/Z and 203139/A/16/Z).

