## Supplementary Materials for "Hemispheric laterality of the putamen predicts pseudoneglect"

| Predictors | AIC | BIC |
|---|---|---|
| ('Puta',) | 33.64 | 37.21 |
| ('Puta', 'Amyg') | 33.80 | 39.15 |
| ('Caud', 'Puta', 'Caud*Puta') | 34.34 | 41.48 |
| ('Puta', 'Pall', 'Pall*Puta') | 35.13 | 42.27 |
| ('Caud', 'Puta', 'Amyg', 'Caud*Puta') | 35.36 | 44.28 |
| ('Puta', 'Hipp') | 35.36 | 40.71 |
| ('Thal', 'Puta', 'Thal*Puta') | 35.44 | 42.57 |
| ('Thal', 'Puta', 'Amyg') | 35.44 | 42.58 |
| ('Thal', 'Puta') | 35.47 | 40.82 |
| ('Caud', 'Puta') | 35.57 | 40.92 |
| ('Puta', 'Hipp', 'Amyg') | 35.61 | 42.75 |
| ('Puta', 'Accu') | 35.62 | 40.97 |
| ('Puta', 'Pall') | 35.64 | 40.99 |
| ('Puta', 'Amyg', 'Accu') | 35.71 | 42.85 |
| ('Puta', 'Pall', 'Amyg') | 35.78 | 42.92 |
| ('Caud', 'Puta', 'Amyg') | 35.80 | 42.93 |
| ('Caud', 'Puta', 'Hipp', 'Caud*Puta') | 36.22 | 45.14 |
| ('Thal', 'Caud', 'Puta', 'Caud*Puta') | 36.28 | 45.20 |
| ('Caud', 'Puta', 'Pall', 'Caud*Puta') | 36.29 | 45.21 |
| ('Caud', 'Puta', 'Accu', 'Caud*Puta') | 36.32 | 45.24 |
| ('Puta', 'Pall', 'Amyg', 'Pall*Puta') | 36.41 | 45.33 |
| ('Thal', 'Puta', 'Amyg', 'Thal*Puta') | 36.52 | 45.44 |
| ('Puta', 'Pall', 'Accu', 'Pall*Puta') | 36.95 | 45.88 |
| ('Puta', 'Pall', 'Hipp', 'Pall*Puta') | 36.99 | 45.91 |
| ('Caud', 'Puta', 'Pall', 'Pall*Puta') | 37.02 | 45.94 |
| ('Thal', 'Puta', 'Pall', 'Pall*Puta') | 37.02 | 45.94 |
| ('Thal', 'Puta', 'Hipp', 'Thal*Puta') | 37.16 | 46.08 |
| ('Thal', 'Caud', 'Puta', 'Thal*Puta') | 37.21 | 46.13 |
| ('Thal', 'Caud', 'Puta', 'Amyg', 'Caud*Puta') | 37.21 | 47.92 |
| ('Thal', 'Puta', 'Accu', 'Thal*Puta') | 37.24 | 46.17 |
| ('Thal', 'Puta', 'Pall', 'Amyg') | 37.25 | 46.17 |
| ('Caud', 'Puta', 'Hipp', 'Amyg', 'Caud*Puta') | 37.26 | 47.97 |
| ('Caud', 'Puta', 'Pall', 'Amyg', 'Caud*Puta') | 37.27 | 47.98 |
| ('Puta', 'Hipp', 'Accu') | 37.29 | 44.43 |
| ('Caud', 'Puta', 'Amyg', 'Accu', 'Caud*Puta') | 37.29 | 48.00 |
| ('Thal', 'Puta', 'Amyg', 'Accu') | 37.30 | 46.22 |

| | | |
|---|---|---|
| **('Thal', 'Puta', 'Pall', 'Thal*Puta')** | 37.30 | 46.22 |
| **('Thal', 'Puta', 'Hipp')** | 37.31 | 44.45 |
| **('Caud', 'Puta', 'Hipp')** | 37.31 | 44.45 |
| **('Puta', 'Pall', 'Hipp')** | 37.33 | 44.46 |
| **('Thal', 'Caud', 'Puta')** | 37.34 | 44.48 |
| **('Thal', 'Puta', 'Pall')** | 37.39 | 44.53 |
| **('Thal', 'Puta', 'Hipp', 'Amyg')** | 37.41 | 46.33 |
| **('Thal', 'Puta', 'Accu')** | 37.42 | 44.56 |
| **('Thal', 'Caud', 'Puta', 'Amyg')** | 37.44 | 46.36 |
| **('Puta', 'Hipp', 'Amyg', 'Accu')** | 37.47 | 46.39 |
| **('Caud', 'Puta', 'Accu')** | 37.54 | 44.68 |
| **('Caud', 'Puta', 'Pall')** | 37.57 | 44.70 |
| **('Puta', 'Pall', 'Hipp', 'Amyg')** | 37.57 | 46.49 |
| **('Caud', 'Puta', 'Hipp', 'Amyg')** | 37.61 | 46.53 |
| **('Puta', 'Pall', 'Accu')** | 37.62 | 44.76 |
| **('Caud', 'Puta', 'Pall', 'Caud*Puta', 'Pall*Puta')** | 37.63 | 48.33 |
| **('Puta', 'Pall', 'Amyg', 'Accu')** | 37.71 | 46.63 |
| **('Caud', 'Puta', 'Amyg', 'Accu')** | 37.71 | 46.63 |
| **('Caud', 'Puta', 'Pall', 'Pall*Puta', 'Caud*Puta*Pall')** | 37.72 | 48.42 |
| **('Caud', 'Puta', 'Pall', 'Amyg')** | 37.78 | 46.70 |
| **('Thal', 'Caud', 'Puta', 'Caud*Puta', 'Thal*Puta')** | 37.88 | 48.58 |
| **('Caud', 'Puta', 'Pall', 'Caud*Puta', 'Caud*Puta*Pall')** | 38.11 | 48.81 |
| **('Thal', 'Puta', 'Pall', 'Amyg', 'Pall*Puta')** | 38.12 | 48.82 |
| **('Caud', 'Puta', 'Pall', 'Hipp', 'Caud*Puta')** | 38.12 | 48.83 |
| **('Thal', 'Caud', 'Puta', 'Pall', 'Caud*Puta')** | 38.14 | 48.84 |
| **('Caud', 'Puta', 'Hipp', 'Accu', 'Caud*Puta')** | 38.18 | 48.88 |
| **('Puta', 'Pall', 'Amyg', 'Accu', 'Pall*Puta')** | 38.20 | 48.90 |
| **('Thal', 'Caud', 'Puta', 'Hipp', 'Caud*Puta')** | 38.21 | 48.91 |
| **('Thal', 'Caud', 'Puta', 'Accu', 'Caud*Puta')** | 38.24 | 48.94 |
| **('Thal',)** | 38.25 | 41.82 |
| **('Caud', 'Puta', 'Pall', 'Caud*Puta', 'Caud*Pall')** | 38.27 | 48.97 |
| **('Thal', 'Puta', 'Amyg', 'Accu', 'Thal*Puta')** | 38.27 | 48.97 |
| **('Caud', 'Puta', 'Pall', 'Accu', 'Caud*Puta')** | 38.28 | 48.98 |

| | | |
|---|---|---|
| **('Puta', 'Pall', 'Hipp', 'Amyg', 'Pall*Puta')** | 38.28 | 48.99 |
| **('Thal', 'Puta', 'Pall', 'Amyg', 'Thal*Puta')** | 38.31 | 49.02 |
| **('Caud', 'Puta', 'Pall', 'Amyg', 'Pall*Puta', 'Caud*Puta*Pall')** | 38.39 | 50.88 |
| **('Caud', 'Puta', 'Pall', 'Amyg', 'Pall*Puta')** | 38.39 | 49.09 |
| **('Thal', 'Puta', 'Hipp', 'Amyg', 'Thal*Puta')** | 38.40 | 49.11 |
| **('Thal', 'Caud', 'Puta', 'Amyg', 'Thal*Puta')** | 38.46 | 49.16 |
| **('Caud', 'Puta', 'Pall', 'Amyg', 'Caud*Puta', 'Caud*Puta*Pall')** | 38.47 | 50.96 |
| **('Puta', 'Pall', 'Hipp', 'Accu', 'Pall*Puta')** | 38.75 | 49.46 |
| **('Caud', 'Puta', 'Pall', 'Amyg', 'Caud*Puta*Pall')** | 38.78 | 49.48 |
| **('Thal', 'Puta', 'Pall', 'Accu', 'Pall*Puta')** | 38.81 | 49.51 |
| **('Caud', 'Puta', 'Pall', 'Accu', 'Pall*Puta')** | 38.82 | 49.53 |
| **('Caud', 'Puta', 'Pall', 'Caud*Puta', 'Pall*Puta', 'Caud*Puta*Pall')** | 38.83 | 51.32 |
| **('Caud', 'Puta', 'Pall', 'Caud*Pall', 'Pall*Puta', 'Caud*Puta*Pall')** | 38.87 | 51.36 |
| **('Thal', 'Puta', 'Hipp', 'Accu', 'Thal*Puta')** | 38.87 | 49.58 |
| **('Thal', 'Caud', 'Puta', 'Pall', 'Pall*Puta')** | 38.88 | 49.58 |
| **('Thal', 'Puta', 'Pall', 'Pall*Puta', 'Thal*Puta')** | 38.88 | 49.59 |
| **('Caud', 'Puta', 'Pall', 'Hipp', 'Pall*Puta')** | 38.91 | 49.62 |
| **('Thal', 'Puta', 'Pall', 'Hipp', 'Pall*Puta')** | 38.94 | 49.65 |
| **('Thal', 'Caud', 'Puta', 'Pall', 'Amyg', 'Caud*Puta')** | 38.96 | 51.44 |
| **('Thal', 'Caud', 'Puta', 'Accu', 'Thal*Puta')** | 38.99 | 49.69 |
| **('Thal', 'Puta', 'Pall', 'Hipp', 'Thal*Puta')** | 39.01 | 49.71 |
| **('Caud', 'Puta', 'Pall', 'Caud*Pall', 'Pall*Puta')** | 39.01 | 49.71 |
| **('Caud', 'Puta', 'Pall', 'Amyg', 'Caud*Puta', 'Pall*Puta')** | 39.01 | 51.50 |
| **('Thal', 'Caud', 'Puta', 'Hipp', 'Thal*Puta')** | 39.02 | 49.72 |
| **('Thal', 'Puta', 'Pall', 'Amyg', 'Accu')** | 39.11 | 49.82 |

| | | |
|---|---|---|
| **('Thal', 'Caud', 'Puta', 'Amyg', 'Accu', 'Caud*Puta')** | 39.11 | 51.60 |
| **('Thal', 'Caud', 'Puta', 'Pall', 'Thal*Puta')** | 39.12 | 49.82 |
| **('Thal', 'Puta', 'Pall', 'Accu', 'Thal*Puta')** | 39.12 | 49.82 |
| **('Caud', 'Puta', 'Pall', 'Hipp', 'Amyg', 'Caud*Puta')** | 39.13 | 51.61 |
| **('Thal', 'Caud', 'Puta', 'Amyg', 'Caud*Puta', 'Thal*Puta')** | 39.13 | 51.62 |
| **('Caud', 'Puta', 'Pall', 'Accu', 'Pall*Puta', 'Caud*Puta*Pall')** | 39.13 | 51.62 |
| **('Caud', 'Puta', 'Pall', 'Amyg', 'Caud*Puta', 'Caud*Pall')** | 39.16 | 51.65 |
| **('Caud', 'Puta', 'Hipp', 'Amyg', 'Accu', 'Caud*Puta')** | 39.17 | 51.65 |
| **('Thal', 'Caud', 'Puta', 'Hipp', 'Amyg', 'Caud*Puta')** | 39.19 | 51.68 |
| **('Thal', 'Amyg')** | 39.21 | 44.56 |
| **('Thal', 'Puta', 'Pall', 'Hipp', 'Amyg')** | 39.21 | 49.92 |
| **('Caud', 'Puta', 'Pall', 'Amyg', 'Accu', 'Caud*Puta')** | 39.22 | 51.71 |
| **('Thal', 'Puta', 'Pall', 'Hipp')** | 39.22 | 48.14 |
| **('Thal', 'Caud', 'Puta', 'Hipp')** | 39.23 | 48.15 |
| **('Thal', 'Puta', 'Hipp', 'Amyg', 'Accu')** | 39.23 | 49.94 |
| **('Thal', 'Puta', 'Hipp', 'Accu')** | 39.23 | 48.16 |
| **('Caud', 'Puta', 'Hipp', 'Accu')** | 39.24 | 48.16 |
| **('Thal', 'Caud', 'Puta', 'Pall', 'Amyg')** | 39.25 | 49.96 |
| **('Puta', 'Pall', 'Hipp', 'Accu')** | 39.27 | 48.19 |
| **('Thal', 'Caud', 'Puta', 'Amyg', 'Accu')** | 39.28 | 49.99 |
| **('Thal', 'Caud', 'Puta', 'Accu')** | 39.29 | 48.21 |
| **('Thal', 'Caud', 'Puta', 'Pall')** | 39.29 | 48.21 |
| **('Caud', 'Puta', 'Pall', 'Hipp')** | 39.29 | 48.22 |
| **('Caud', 'Puta', 'Pall', 'Caud*Pall')** | 39.33 | 48.26 |
| **('Caud', 'Puta', 'Pall', 'Amyg', 'Caud*Pall')** | 39.35 | 50.06 |
| **('Thal', 'Puta', 'Pall', 'Accu')** | 39.35 | 48.27 |
| **('Amyg',)** | 39.36 | 42.92 |
| **('Caud', 'Puta', 'Pall', 'Caud*Puta*Pall')** | 39.39 | 48.31 |

| | | |
|---|---|---|
| **('Thal', 'Caud', 'Puta', 'Hipp', 'Amyg')** | 39.40 | 50.11 |
| **('Puta', 'Pall', 'Hipp', 'Amyg', 'Accu')** | 39.44 | 50.14 |
| **('Caud', 'Puta', 'Hipp', 'Amyg', 'Accu')** | 39.47 | 50.17 |
| **('Thal', 'Caud', 'Puta', 'Pall', 'Caud*Puta', 'Pall*Puta')** | 39.52 | 52.01 |
| **('Caud', 'Puta', 'Pall', 'Hipp', 'Caud*Puta', 'Pall*Puta')** | 39.53 | 52.01 |
| **('Caud', 'Puta', 'Pall', 'Accu', 'Caud*Puta', 'Pall*Puta')** | 39.54 | 52.03 |
| **('Caud', 'Puta', 'Pall', 'Accu')** | 39.54 | 48.46 |
| **('Caud', 'Puta', 'Pall', 'Hipp', 'Amyg')** | 39.56 | 50.26 |
| **('Hipp',)** | 39.59 | 43.16 |
| **('Caud', 'Puta', 'Pall', 'Amyg', 'Accu', 'Pall*Puta', 'Caud*Puta*Pall')** | 39.60 | 53.87 |
| **('Caud', 'Puta', 'Pall', 'Caud*Puta', 'Caud*Pall', 'Pall*Puta')** | 39.62 | 52.11 |
| **('Caud', 'Puta', 'Pall', 'Amyg', 'Caud*Puta', 'Pall*Puta', 'Caud*Puta*Pall')** | 39.66 | 53.93 |
| **('Caud', 'Puta', 'Pall', 'Hipp', 'Pall*Puta', 'Caud*Puta*Pall')** | 39.68 | 52.17 |
| **('Caud', 'Puta', 'Pall', 'Amyg', 'Accu')** | 39.71 | 50.41 |
| **('Thal', 'Caud', 'Puta', 'Pall', 'Pall*Puta', 'Caud*Puta*Pall')** | 39.71 | 52.20 |
| **('Accu',)** | 39.74 | 43.31 |
| **('Thal', 'Caud', 'Puta', 'Pall', 'Caud*Puta', 'Thal*Puta')** | 39.74 | 52.23 |
| **('Thal', 'Caud', 'Puta', 'Hipp', 'Caud*Puta', 'Thal*Puta')** | 39.77 | 52.26 |
| **('Thal', 'Caud', 'Puta', 'Accu', 'Caud*Puta', 'Thal*Puta')** | 39.77 | 52.26 |
| **('Pall',)** | 39.77 | 43.34 |
| **('Thal', 'Puta', 'Pall', 'Amyg', 'Accu', 'Pall*Puta')** | 39.85 | 52.34 |
| **('Caud', 'Puta', 'Pall', 'Amyg', 'Caud*Pall', 'Pall*Puta', 'Caud*Puta*Pall')** | 39.87 | 54.15 |
| **('Caud',)** | 39.91 | 43.48 |
| **('Thal', 'Accu')** | 39.95 | 45.30 |

| | | |
|---|---|---|
| **('Caud', 'Puta', 'Pall', 'Hipp', 'Caud*Puta', 'Caud*Puta*Pall')** | 39.96 | 52.45 |
| **('Puta', 'Pall', 'Hipp', 'Amyg', 'Accu', 'Pall*Puta')** | 40.01 | 52.50 |
| **('Thal', 'Caud', 'Puta', 'Pall', 'Caud*Puta', 'Caud*Puta*Pall')** | 40.03 | 52.52 |
| **('Thal', 'Puta', 'Pall', 'Amyg', 'Pall*Puta', 'Thal*Puta')** | 40.06 | 52.54 |
| **('Thal', 'Caud', 'Puta', 'Pall', 'Hipp', 'Caud*Puta')** | 40.06 | 52.55 |
| **('Thal', 'Puta', 'Pall', 'Amyg', 'Accu', 'Thal*Puta')** | 40.07 | 52.56 |
| **('Caud', 'Puta', 'Pall', 'Accu', 'Caud*Puta', 'Caud*Puta*Pall')** | 40.07 | 52.56 |
| **('Thal', 'Puta', 'Hipp', 'Amyg', 'Accu', 'Thal*Puta')** | 40.08 | 52.57 |
| **('Caud', 'Puta', 'Pall', 'Hipp', 'Accu', 'Caud*Puta')** | 40.09 | 52.58 |
| **('Thal', 'Puta', 'Pall', 'Hipp', 'Amyg', 'Pall*Puta')** | 40.09 | 52.58 |
| **('Thal', 'Caud', 'Puta', 'Pall', 'Amyg', 'Pall*Puta')** | 40.10 | 52.58 |
| **('Caud', 'Puta', 'Pall', 'Hipp', 'Caud*Puta', 'Caud*Pall')** | 40.11 | 52.59 |
| **('Caud', 'Puta', 'Pall', 'Caud*Puta', 'Caud*Pall', 'Caud*Puta*Pall')** | 40.11 | 52.60 |
| **('Thal', 'Caud', 'Puta', 'Pall', 'Accu', 'Caud*Puta')** | 40.11 | 52.60 |
| **('Thal', 'Caud', 'Puta', 'Pall', 'Caud*Puta', 'Caud*Pall')** | 40.13 | 52.62 |
| **('Thal', 'Caud', 'Puta', 'Hipp', 'Accu', 'Caud*Puta')** | 40.16 | 52.65 |
| **('Caud', 'Puta', 'Pall', 'Amyg', 'Accu', 'Pall*Puta')** | 40.17 | 52.66 |
| **('Thal', 'Pall')** | 40.17 | 45.53 |
| **('Thal', 'Puta', 'Pall', 'Hipp', 'Amyg', 'Thal*Puta')** | 40.18 | 52.67 |
| **('Thal', 'Caud', 'Puta', 'Amyg', 'Accu', 'Thal*Puta')** | 40.19 | 52.68 |
| **('Thal', 'Caud')** | 40.20 | 45.55 |
| **('Caud', 'Puta', 'Pall', 'Caud*Puta', 'Caud*Pall', 'Pall*Puta', 'Caud*Puta*Pall')** | 40.23 | 54.50 |
| **('Caud', 'Puta', 'Pall', 'Accu', 'Caud*Puta', 'Caud*Pall')** | 40.23 | 52.72 |
| **('Thal', 'Hipp')** | 40.25 | 45.60 |

| | | |
|---|---|---|
| **('Thal', 'Caud', 'Puta', 'Pall', 'Amyg', 'Thal*Puta')** | 40.28 | 52.77 |
| **('Caud', 'Puta', 'Pall', 'Hipp', 'Amyg', 'Pall*Puta')** | 40.28 | 52.77 |
| **('Caud', 'Puta', 'Pall', 'Amyg', 'Accu', 'Caud*Puta', 'Caud*Puta*Pall')** | 40.30 | 54.57 |
| **('Thal', 'Caud', 'Puta', 'Pall', 'Amyg', 'Caud*Puta', 'Caud*Puta*Pall')** | 40.35 | 54.63 |
| **('Thal', 'Caud', 'Puta', 'Hipp', 'Amyg', 'Thal*Puta')** | 40.36 | 52.85 |
| **('Caud', 'Puta', 'Pall', 'Hipp', 'Amyg', 'Pall*Puta', 'Caud*Puta*Pall')** | 40.36 | 54.64 |
| **('Caud', 'Puta', 'Pall', 'Amyg', 'Caud*Pall', 'Pall*Puta')** | 40.37 | 52.86 |
| **('Caud', 'Puta', 'Pall', 'Hipp', 'Amyg', 'Caud*Puta', 'Caud*Puta*Pall')** | 40.38 | 54.65 |
| **('Thal', 'Caud', 'Puta', 'Pall', 'Amyg', 'Pall*Puta', 'Caud*Puta*Pall')** | 40.38 | 54.65 |
| **('Thal', 'Caud', 'Puta', 'Pall', 'Thal*Puta', 'Caud*Puta*Pall')** | 40.46 | 52.95 |
| **('Caud', 'Puta', 'Pall', 'Amyg', 'Caud*Puta', 'Caud*Pall', 'Caud*Puta*Pall')** | 40.47 | 54.75 |
| **('Caud', 'Puta', 'Pall', 'Accu', 'Caud*Puta', 'Pall*Puta', 'Caud*Puta*Pall')** | 40.48 | 54.76 |
| **('Caud', 'Puta', 'Pall', 'Accu', 'Caud*Pall', 'Pall*Puta', 'Caud*Puta*Pall')** | 40.52 | 54.79 |
| **('Caud', 'Puta', 'Pall', 'Amyg', 'Accu', 'Caud*Puta*Pall')** | 40.52 | 53.01 |
| **('Thal', 'Caud', 'Puta', 'Pall', 'Amyg', 'Caud*Puta*Pall')** | 40.54 | 53.03 |
| **('Caud', 'Puta', 'Pall', 'Hipp', 'Amyg', 'Caud*Puta*Pall')** | 40.62 | 53.11 |
| **('Thal', 'Caud', 'Puta', 'Pall', 'Accu', 'Pall*Puta')** | 40.64 | 53.13 |
| **('Caud', 'Puta', 'Pall', 'Amyg', 'Caud*Pall', 'Caud*Puta*Pall')** | 40.65 | 53.14 |
| **('Thal', 'Puta', 'Pall', 'Accu', 'Pall*Puta', 'Thal*Puta')** | 40.66 | 53.15 |

| | | |
|---|---|---|
| **('Caud', 'Puta', 'Pall', 'Hipp', 'Accu', 'Pall*Puta')** | 40.66 | 53.15 |
| **('Thal', 'Puta', 'Pall', 'Hipp', 'Accu', 'Pall*Puta')** | 40.69 | 53.18 |
| **('Thal', 'Amyg', 'Accu')** | 40.71 | 47.85 |
| **('Thal', 'Caud', 'Puta', 'Pall', 'Pall*Puta', 'Thal*Puta')** | 40.72 | 53.21 |
| **('Thal', 'Caud', 'Puta', 'Hipp', 'Accu', 'Thal*Puta')** | 40.72 | 53.21 |
| **('Thal', 'Puta', 'Pall', 'Hipp', 'Pall*Puta', 'Thal*Puta')** | 40.73 | 53.22 |
| **('Thal', 'Puta', 'Pall', 'Hipp', 'Accu', 'Thal*Puta')** | 40.73 | 53.22 |
| **('Thal', 'Caud', 'Puta', 'Pall', 'Amyg', 'Caud*Puta', 'Pall*Puta')** | 40.76 | 55.04 |
| **('Caud', 'Puta', 'Pall', 'Hipp', 'Caud*Puta', 'Pall*Puta', 'Caud*Puta*Pall')** | 40.79 | 55.06 |
| **('Caud', 'Puta', 'Pall', 'Accu', 'Caud*Pall', 'Pall*Puta')** | 40.82 | 53.31 |
| **('Thal', 'Caud', 'Puta', 'Pall', 'Caud*Puta', 'Pall*Puta', 'Caud*Puta*Pall')** | 40.83 | 55.11 |
| **('Thal', 'Caud', 'Puta', 'Pall', 'Hipp', 'Pall*Puta')** | 40.84 | 53.33 |
| **('Thal', 'Caud', 'Puta', 'Pall', 'Caud*Pall', 'Pall*Puta', 'Caud*Puta*Pall')** | 40.85 | 55.12 |
| **('Thal', 'Caud', 'Puta', 'Pall', 'Caud*Pall', 'Pall*Puta')** | 40.86 | 53.35 |
| **('Caud', 'Puta', 'Pall', 'Hipp', 'Caud*Pall', 'Pall*Puta', 'Caud*Puta*Pall')** | 40.86 | 55.13 |
| **('Thal', 'Caud', 'Puta', 'Pall', 'Amyg', 'Accu', 'Caud*Puta')** | 40.87 | 55.14 |
| **('Thal', 'Caud', 'Puta', 'Pall', 'Amyg', 'Caud*Puta', 'Caud*Pall')** | 40.88 | 55.15 |
| **('Thal', 'Caud', 'Puta', 'Pall', 'Amyg', 'Caud*Puta', 'Thal*Puta')** | 40.88 | 55.16 |
| **('Caud', 'Puta', 'Pall', 'Hipp', 'Amyg', 'Caud*Puta', 'Pall*Puta')** | 40.90 | 55.18 |

| | | |
|---|---|---|
| **('Caud', 'Puta', 'Pall', 'Hipp', 'Caud*Pall', 'Pall*Puta')** | 40.90 | 53.39 |
| **('Thal', 'Caud', 'Puta', 'Pall', 'Accu', 'Thal*Puta')** | 40.91 | 53.40 |
| **('Thal', 'Caud', 'Puta', 'Pall', 'Hipp', 'Thal*Puta')** | 40.91 | 53.40 |
| **('Caud', 'Puta', 'Pall', 'Amyg', 'Accu', 'Caud*Puta', 'Pall*Puta')** | 40.91 | 55.18 |
| **('Thal', 'Caud', 'Puta', 'Pall', 'Amyg', 'Caud*Pall')** | 40.92 | 53.41 |
| **('Thal', 'Caud', 'Puta', 'Pall', 'Hipp', 'Amyg', 'Caud*Puta')** | 40.92 | 55.20 |
| **('Thal', 'Caud', 'Puta', 'Amyg', 'Accu', 'Caud*Puta', 'Thal*Puta')** | 40.99 | 55.26 |
| **('Caud', 'Puta', 'Pall', 'Amyg', 'Caud*Puta', 'Caud*Pall', 'Pall*Puta')** | 40.99 | 55.27 |
| **('Caud', 'Puta', 'Pall', 'Amyg', 'Accu', 'Caud*Puta', 'Caud*Pall')** | 41.02 | 55.30 |
| **('Caud', 'Puta', 'Pall', 'Hipp', 'Amyg', 'Caud*Puta', 'Caud*Pall')** | 41.03 | 55.30 |
| **('Caud', 'Puta', 'Pall', 'Hipp', 'Amyg', 'Accu', 'Caud*Puta')** | 41.04 | 55.32 |
| **('Thal', 'Puta', 'Pall', 'Hipp', 'Amyg', 'Accu')** | 41.05 | 53.54 |
| **('Thal', 'Pall', 'Amyg')** | 41.05 | 48.19 |
| **('Caud', 'Puta', 'Pall', 'Amyg', 'Accu', 'Caud*Pall')** | 41.05 | 53.54 |
| **('Caud', 'Puta', 'Pall', 'Hipp', 'Accu', 'Pall*Puta', 'Caud*Puta*Pall')** | 41.06 | 55.33 |
| **('Thal', 'Caud', 'Puta', 'Hipp', 'Amyg', 'Accu', 'Caud*Puta')** | 41.07 | 55.35 |
| **('Amyg', 'Accu')** | 41.08 | 46.43 |
| **('Thal', 'Caud', 'Puta', 'Pall', 'Amyg', 'Thal*Puta', 'Caud*Puta*Pall')** | 41.08 | 55.36 |
| **('Thal', 'Caud', 'Puta', 'Hipp', 'Amyg', 'Caud*Puta', 'Thal*Puta')** | 41.09 | 55.36 |
| **('Hipp', 'Amyg')** | 41.09 | 46.44 |
| **('Caud', 'Puta', 'Pall', 'Hipp', 'Caud*Pall')** | 41.09 | 51.80 |

| | | |
|---|---|---|
| **('Thal', 'Caud', 'Puta', 'Pall', 'Caud*Pall', 'Thal*Puta')** | 41.10 | 53.59 |
| **('Thal', 'Caud', 'Puta', 'Pall', 'Amyg', 'Accu')** | 41.11 | 53.60 |
| **('Thal', 'Caud', 'Puta', 'Pall', 'Caud*Pall')** | 41.12 | 51.82 |
| **('Caud', 'Puta', 'Pall', 'Amyg', 'Accu', 'Caud*Puta', 'Pall*Puta', 'Caud*Puta*Pall')** | 41.13 | 57.18 |
| **('Thal', 'Caud', 'Puta', 'Pall', 'Accu', 'Pall*Puta', 'Caud*Puta*Pall')** | 41.13 | 55.41 |
| **('Caud', 'Puta', 'Pall', 'Hipp', 'Caud*Puta*Pall')** | 41.15 | 51.85 |
| **('Thal', 'Caud', 'Puta', 'Hipp', 'Accu')** | 41.15 | 51.86 |
| **('Thal', 'Puta', 'Pall', 'Hipp', 'Accu')** | 41.16 | 51.86 |
| **('Caud', 'Puta', 'Pall', 'Hipp', 'Amyg', 'Caud*Pall')** | 41.16 | 53.65 |
| **('Thal', 'Caud', 'Puta', 'Pall', 'Hipp')** | 41.17 | 51.87 |
| **('Thal', 'Hipp', 'Amyg')** | 41.20 | 48.33 |
| **('Thal', 'Caud', 'Amyg')** | 41.21 | 48.34 |
| **('Thal', 'Caud', 'Puta', 'Pall', 'Hipp', 'Amyg')** | 41.21 | 53.70 |
| **('Thal', 'Caud', 'Puta', 'Pall', 'Caud*Puta*Pall')** | 41.21 | 51.92 |
| **('Pall', 'Amyg')** | 41.21 | 46.56 |
| **('Caud', 'Puta', 'Pall', 'Accu', 'Caud*Pall')** | 41.22 | 51.92 |
| **('Thal', 'Caud', 'Puta', 'Hipp', 'Amyg', 'Accu')** | 41.23 | 53.72 |
| **('Caud', 'Puta', 'Pall', 'Hipp', 'Accu')** | 41.23 | 51.94 |
| **('Thal', 'Caud', 'Puta', 'Pall', 'Accu')** | 41.24 | 51.94 |
| **('Caud', 'Puta', 'Pall', 'Caud*Pall', 'Caud*Puta*Pall')** | 41.27 | 51.97 |
| **('Hipp', 'Accu')** | 41.29 | 46.65 |
| **('Caud', 'Amyg')** | 41.29 | 46.65 |
| **('Caud', 'Puta', 'Pall', 'Amyg', 'Caud*Puta', 'Caud*Pall', 'Pall*Puta', 'Caud*Puta*Pall')** | 41.30 | 57.36 |
| **('Caud', 'Puta', 'Pall', 'Amyg', 'Accu', 'Caud*Pall', 'Pall*Puta', 'Caud*Puta*Pall')** | 41.31 | 57.37 |
| **('Caud', 'Puta', 'Pall', 'Accu', 'Caud*Puta*Pall')** | 41.34 | 52.05 |

| | | |
|---|---|---|
| **('Thal', 'Caud', 'Puta', 'Pall', 'Caud*Puta', 'Thal*Puta', 'Caud*Puta*Pall')** | 41.36 | 55.64 |
| **('Caud', 'Puta', 'Pall', 'Hipp', 'Accu', 'Caud*Puta', 'Pall*Puta')** | 41.40 | 55.67 |
| **('Thal', 'Caud', 'Puta', 'Pall', 'Accu', 'Caud*Puta', 'Pall*Puta')** | 41.40 | 55.67 |
| **('Caud', 'Puta', 'Pall', 'Hipp', 'Amyg', 'Accu')** | 41.43 | 53.92 |
| **('Thal', 'Caud', 'Puta', 'Pall', 'Hipp', 'Caud*Puta', 'Pall*Puta')** | 41.47 | 55.75 |
| **('Thal', 'Caud', 'Puta', 'Pall', 'Caud*Puta', 'Caud*Pall', 'Pall*Puta')** | 41.50 | 55.77 |
| **('Thal', 'Caud', 'Puta', 'Pall', 'Caud*Puta', 'Pall*Puta', 'Thal*Puta')** | 41.51 | 55.78 |
| **('Caud', 'Puta', 'Pall', 'Hipp', 'Caud*Puta', 'Caud*Pall', 'Pall*Puta')** | 41.51 | 55.79 |
| **('Pall', 'Hipp')** | 41.52 | 46.87 |
| **('Caud', 'Puta', 'Pall', 'Hipp', 'Amyg', 'Accu', 'Pall*Puta', 'Caud*Puta*Pall')** | 41.53 | 57.59 |
| **('Caud', 'Puta', 'Pall', 'Accu', 'Caud*Puta', 'Caud*Pall', 'Pall*Puta')** | 41.54 | 55.81 |
| **('Pall', 'Accu')** | 41.56 | 46.92 |
| **('Caud', 'Hipp')** | 41.57 | 46.93 |
| **('Thal', 'Caud', 'Puta', 'Pall', 'Pall*Puta', 'Thal*Puta', 'Caud*Puta*Pall')** | 41.58 | 55.85 |
| **('Thal', 'Caud', 'Puta', 'Pall', 'Amyg', 'Accu', 'Pall*Puta', 'Caud*Puta*Pall')** | 41.59 | 57.65 |
| **('Thal', 'Caud', 'Puta', 'Pall', 'Hipp', 'Caud*Puta', 'Thal*Puta')** | 41.61 | 55.89 |
| **('Thal', 'Caud', 'Puta', 'Hipp', 'Accu', 'Caud*Puta', 'Thal*Puta')** | 41.62 | 55.89 |
| **('Caud', 'Puta', 'Pall', 'Hipp', 'Amyg', 'Caud*Puta', 'Pall*Puta', 'Caud*Puta*Pall')** | 41.62 | 57.68 |

| | | |
|---|---|---|
| **('Thal', 'Caud', 'Puta', 'Pall', 'Amyg', 'Caud*Puta', 'Pall*Puta', 'Caud*Puta*Pall')** | 41.64 | 57.70 |
| **('Thal', 'Caud', 'Puta', 'Pall', 'Accu', 'Caud*Puta', 'Thal*Puta')** | 41.64 | 55.92 |
| **('Thal', 'Caud', 'Puta', 'Pall', 'Hipp', 'Pall*Puta', 'Caud*Puta*Pall')** | 41.67 | 55.94 |
| **('Thal', 'Caud', 'Puta', 'Pall', 'Caud*Puta', 'Caud*Pall', 'Thal*Puta')** | 41.73 | 56.00 |
| **('Caud', 'Accu')** | 41.73 | 47.09 |
| **('Caud', 'Pall')** | 41.77 | 47.13 |
| **('Thal', 'Puta', 'Pall', 'Amyg', 'Accu', 'Pall*Puta', 'Thal*Puta')** | 41.78 | 56.05 |
| **('Thal', 'Puta', 'Pall', 'Hipp', 'Amyg', 'Accu', 'Pall*Puta')** | 41.80 | 56.07 |
| **('Thal', 'Caud', 'Puta', 'Pall', 'Amyg', 'Accu', 'Pall*Puta')** | 41.82 | 56.09 |
| **('Caud', 'Puta', 'Pall', 'Hipp', 'Amyg', 'Caud*Pall', 'Pall*Puta', 'Caud*Puta*Pall')** | 41.86 | 57.92 |
| **('Thal', 'Caud', 'Puta', 'Pall', 'Amyg', 'Caud*Pall', 'Pall*Puta', 'Caud*Puta*Pall')** | 41.87 | 57.93 |
| **('Thal', 'Puta', 'Pall', 'Hipp', 'Amyg', 'Accu', 'Thal*Puta')** | 41.87 | 56.15 |
| **('Thal', 'Pall', 'Accu')** | 41.88 | 49.02 |
| **('Thal', 'Caud', 'Accu')** | 41.88 | 49.02 |
| **('Caud', 'Puta', 'Pall', 'Hipp', 'Accu', 'Caud*Puta', 'Caud*Puta*Pall')** | 41.89 | 56.17 |
| **('Thal', 'Hipp', 'Accu')** | 41.92 | 49.06 |
| **('Thal', 'Caud', 'Puta', 'Pall', 'Hipp', 'Caud*Puta', 'Caud*Puta*Pall')** | 41.94 | 56.21 |
| **('Caud', 'Puta', 'Pall', 'Hipp', 'Caud*Puta', 'Caud*Pall', 'Caud*Puta*Pall')** | 41.96 | 56.23 |
| **('Thal', 'Caud', 'Puta', 'Pall', 'Accu', 'Caud*Puta', 'Caud*Puta*Pall')** | 41.98 | 56.26 |
| **('Thal', 'Puta', 'Pall', 'Hipp', 'Amyg', 'Pall*Puta', 'Thal*Puta')** | 42.00 | 56.27 |

| | | |
|---|---|---|
| **('Caud', 'Puta', 'Pall', 'Hipp', 'Amyg', 'Accu', 'Pall*Puta')** | 42.01 | 56.28 |
| **('Caud', 'Puta', 'Pall', 'Accu', 'Caud*Puta', 'Caud*Pall', 'Pall*Puta', 'Caud*Puta*Pall')** | 42.01 | 58.07 |
| **('Thal', 'Caud', 'Puta', 'Pall', 'Hipp', 'Accu', 'Caud*Puta')** | 42.02 | 56.29 |
| **('Thal', 'Caud', 'Puta', 'Pall', 'Amyg', 'Accu', 'Thal*Puta')** | 42.02 | 56.29 |
| **('Thal', 'Caud', 'Puta', 'Pall', 'Amyg', 'Pall*Puta', 'Thal*Puta')** | 42.02 | 56.30 |
| **('Thal', 'Caud', 'Puta', 'Pall', 'Caud*Puta', 'Caud*Pall', 'Caud*Puta*Pall')** | 42.03 | 56.30 |
| **('Thal', 'Caud', 'Puta', 'Hipp', 'Amyg', 'Accu', 'Thal*Puta')** | 42.03 | 56.31 |
| **('Caud', 'Puta', 'Pall', 'Hipp', 'Accu', 'Caud*Puta', 'Caud*Pall')** | 42.04 | 56.31 |
| **('Thal', 'Caud', 'Puta', 'Pall', 'Hipp', 'Caud*Puta', 'Caud*Pall')** | 42.05 | 56.32 |
| **('Thal', 'Caud', 'Puta', 'Pall', 'Accu', 'Thal*Puta', 'Caud*Puta*Pall')** | 42.05 | 56.33 |
| **('Caud', 'Puta', 'Pall', 'Accu', 'Caud*Puta', 'Caud*Pall', 'Caud*Puta*Pall')** | 42.06 | 56.34 |
| **('Caud', 'Puta', 'Pall', 'Amyg', 'Accu', 'Caud*Pall', 'Pall*Puta')** | 42.07 | 56.34 |
| **('Thal', 'Caud', 'Puta', 'Pall', 'Hipp', 'Amyg', 'Pall*Puta')** | 42.08 | 56.35 |
| **('Thal', 'Caud', 'Puta', 'Pall', 'Accu', 'Caud*Puta', 'Caud*Pall')** | 42.08 | 56.35 |
| **('Thal', 'Caud', 'Puta', 'Pall', 'Amyg', 'Caud*Pall', 'Pall*Puta')** | 42.08 | 56.35 |
| **('Thal', 'Caud', 'Puta', 'Pall', 'Caud*Pall', 'Thal*Puta', 'Caud*Puta*Pall')** | 42.08 | 56.36 |
| **('Thal', 'Caud', 'Puta', 'Pall', 'Amyg', 'Caud*Puta', 'Thal*Puta', 'Caud*Puta*Pall')** | 42.09 | 58.15 |
| **('Thal', 'Caud', 'Pall')** | 42.14 | 49.28 |

| | | |
|---|---|---|
| **('Thal', 'Caud', 'Puta', 'Pall', 'Amyg', 'Accu', 'Caud*Puta', 'Caud*Puta*Pall')** | 42.15 | 58.20 |
| **('Caud', 'Puta', 'Pall', 'Hipp', 'Amyg', 'Accu', 'Caud*Puta', 'Caud*Puta*Pall')** | 42.15 | 58.21 |
| **('Thal', 'Pall', 'Hipp')** | 42.17 | 49.30 |
| **('Thal', 'Caud', 'Puta', 'Pall', 'Hipp', 'Amyg', 'Thal*Puta')** | 42.17 | 56.44 |
| **('Thal', 'Caud', 'Puta', 'Pall', 'Hipp', 'Thal*Puta', 'Caud*Puta*Pall')** | 42.18 | 56.45 |
| **('Thal', 'Caud', 'Hipp')** | 42.20 | 49.34 |
| **('Caud', 'Puta', 'Pall', 'Hipp', 'Caud*Puta', 'Caud*Pall', 'Pall*Puta', 'Caud*Puta*Pall')** | 42.20 | 58.26 |
| **('Thal', 'Caud', 'Puta', 'Pall', 'Caud*Puta', 'Caud*Pall', 'Pall*Puta', 'Caud*Puta*Pall')** | 42.22 | 58.28 |
| **('Thal', 'Caud', 'Puta', 'Pall', 'Amyg', 'Accu', 'Caud*Puta*Pall')** | 42.24 | 56.52 |
| **('Caud', 'Puta', 'Pall', 'Amyg', 'Accu', 'Caud*Pall', 'Caud*Puta*Pall')** | 42.24 | 56.52 |
| **('Thal', 'Caud', 'Puta', 'Pall', 'Amyg', 'Caud*Pall', 'Thal*Puta')** | 42.25 | 56.53 |
| **('Caud', 'Puta', 'Pall', 'Hipp', 'Amyg', 'Caud*Pall', 'Pall*Puta')** | 42.26 | 56.53 |
| **('Caud', 'Puta', 'Pall', 'Amyg', 'Accu', 'Caud*Puta', 'Caud*Pall', 'Caud*Puta*Pall')** | 42.26 | 58.31 |
| **('Caud', 'Puta', 'Pall', 'Hipp', 'Amyg', 'Accu', 'Caud*Puta*Pall')** | 42.30 | 56.57 |
| **('Thal', 'Caud', 'Puta', 'Pall', 'Hipp', 'Amyg', 'Caud*Puta', 'Caud*Puta*Pall')** | 42.32 | 58.37 |
| **('Thal', 'Caud', 'Puta', 'Pall', 'Amyg', 'Pall*Puta', 'Thal*Puta', 'Caud*Puta*Pall')** | 42.35 | 58.40 |
| **('Thal', 'Caud', 'Puta', 'Pall', 'Amyg', 'Caud*Puta', 'Caud*Pall', 'Caud*Puta*Pall')** | 42.35 | 58.41 |
| **('Thal', 'Caud', 'Puta', 'Pall', 'Hipp', 'Amyg', 'Pall*Puta', 'Caud*Puta*Pall')** | 42.36 | 58.42 |

| | | |
|---|---|---|
| **('Caud', 'Puta', 'Pall', 'Hipp', 'Amyg', 'Caud*Puta', 'Caud*Pall', 'Caud*Puta*Pall')** | 42.37 | 58.43 |
| **('Caud', 'Puta', 'Pall', 'Hipp', 'Accu', 'Caud*Puta', 'Pall*Puta', 'Caud*Puta*Pall')** | 42.40 | 58.46 |
| **('Thal', 'Caud', 'Puta', 'Pall', 'Amyg', 'Caud*Pall', 'Caud*Puta*Pall')** | 42.43 | 56.70 |
| **('Thal', 'Puta', 'Pall', 'Hipp', 'Accu', 'Pall*Puta', 'Thal*Puta')** | 42.44 | 56.71 |
| **('Thal', 'Caud', 'Puta', 'Pall', 'Accu', 'Pall*Puta', 'Thal*Puta')** | 42.46 | 56.74 |
| **('Caud', 'Puta', 'Pall', 'Hipp', 'Accu', 'Caud*Pall', 'Pall*Puta', 'Caud*Puta*Pall')** | 42.47 | 58.53 |
| **('Thal', 'Caud', 'Puta', 'Pall', 'Accu', 'Caud*Puta', 'Pall*Puta', 'Caud*Puta*Pall')** | 42.48 | 58.54 |
| **('Thal', 'Caud', 'Puta', 'Pall', 'Hipp', 'Amyg', 'Caud*Puta*Pall')** | 42.49 | 56.76 |
| **('Thal', 'Caud', 'Puta', 'Pall', 'Amyg', 'Accu', 'Thal*Puta', 'Caud*Puta*Pall')** | 42.50 | 58.55 |
| **('Thal', 'Caud', 'Puta', 'Pall', 'Accu', 'Caud*Pall', 'Pall*Puta', 'Caud*Puta*Pall')** | 42.50 | 58.56 |
| **('Caud', 'Puta', 'Pall', 'Hipp', 'Amyg', 'Caud*Pall', 'Caud*Puta*Pall')** | 42.51 | 56.79 |
| **('Thal', 'Caud', 'Puta', 'Pall', 'Amyg', 'Accu', 'Caud*Pall')** | 42.55 | 56.83 |
| **('Thal', 'Pall', 'Amyg', 'Accu')** | 42.57 | 51.49 |
| **('Thal', 'Caud', 'Puta', 'Pall', 'Hipp', 'Accu', 'Pall*Puta')** | 42.58 | 56.85 |
| **('Thal', 'Caud', 'Puta', 'Pall', 'Caud*Pall', 'Pall*Puta', 'Thal*Puta', 'Caud*Puta*Pall')** | 42.60 | 58.65 |
| **('Thal', 'Caud', 'Puta', 'Pall', 'Amyg', 'Accu', 'Caud*Puta', 'Pall*Puta')** | 42.62 | 58.67 |
| **('Thal', 'Caud', 'Puta', 'Pall', 'Hipp', 'Accu', 'Thal*Puta')** | 42.62 | 56.89 |

| | | |
|---|---|---|
| **('Thal', 'Caud', 'Puta', 'Pall', 'Hipp', 'Pall*Puta', 'Thal*Puta')** | 42.62 | 56.90 |
| **('Thal', 'Caud', 'Puta', 'Pall', 'Accu', 'Caud*Pall', 'Pall*Puta')** | 42.64 | 56.92 |
| **('Caud', 'Puta', 'Pall', 'Hipp', 'Accu', 'Caud*Pall', 'Pall*Puta')** | 42.66 | 56.93 |
| **('Thal', 'Caud', 'Puta', 'Pall', 'Caud*Pall', 'Pall*Puta', 'Thal*Puta')** | 42.68 | 56.95 |
| **('Hipp', 'Amyg', 'Accu')** | 42.69 | 49.83 |
| **('Thal', 'Caud', 'Puta', 'Pall', 'Amyg', 'Accu', 'Caud*Puta', 'Caud*Pall')** | 42.70 | 58.76 |
| **('Thal', 'Caud', 'Amyg', 'Accu')** | 42.71 | 51.63 |
| **('Thal', 'Hipp', 'Amyg', 'Accu')** | 42.71 | 51.63 |
| **('Thal', 'Caud', 'Puta', 'Pall', 'Hipp', 'Amyg', 'Caud*Puta', 'Pall*Puta')** | 42.74 | 58.80 |
| **('Thal', 'Caud', 'Puta', 'Pall', 'Amyg', 'Caud*Puta', 'Caud*Pall', 'Pall*Puta')** | 42.75 | 58.81 |
| **('Thal', 'Caud', 'Puta', 'Pall', 'Amyg', 'Accu', 'Caud*Puta', 'Thal*Puta')** | 42.76 | 58.82 |
| **('Caud', 'Puta', 'Pall', 'Hipp', 'Amyg', 'Accu', 'Caud*Puta', 'Pall*Puta')** | 42.76 | 58.82 |
| **('Thal', 'Caud', 'Puta', 'Pall', 'Amyg', 'Caud*Puta', 'Pall*Puta', 'Thal*Puta')** | 42.76 | 58.82 |
| **('Caud', 'Puta', 'Pall', 'Hipp', 'Amyg', 'Accu', 'Caud*Pall')** | 42.78 | 57.05 |
| **('Thal', 'Caud', 'Puta', 'Pall', 'Hipp', 'Caud*Puta', 'Pall*Puta', 'Caud*Puta*Pall')** | 42.78 | 58.84 |
| **('Thal', 'Caud', 'Puta', 'Pall', 'Hipp', 'Caud*Pall', 'Pall*Puta', 'Caud*Puta*Pall')** | 42.81 | 58.87 |
| **('Thal', 'Caud', 'Puta', 'Pall', 'Caud*Puta', 'Pall*Puta', 'Thal*Puta', 'Caud*Puta*Pall')** | 42.82 | 58.88 |

| | | |
|---|---|---|
| **('Thal', 'Caud', 'Puta', 'Pall', 'Hipp', 'Amyg', 'Accu', 'Caud*Puta')** | 42.82 | 58.88 |
| **('Thal', 'Caud', 'Puta', 'Pall', 'Hipp', 'Caud*Pall', 'Pall*Puta')** | 42.82 | 57.09 |
| **('Thal', 'Caud', 'Puta', 'Pall', 'Hipp', 'Amyg', 'Caud*Puta', 'Thal*Puta')** | 42.83 | 58.89 |
| **('Caud', 'Puta', 'Pall', 'Hipp', 'Amyg', 'Accu', 'Caud*Puta', 'Caud*Pall')** | 42.84 | 58.90 |
| **('Caud', 'Puta', 'Pall', 'Amyg', 'Accu', 'Caud*Puta', 'Caud*Pall', 'Pall*Puta')** | 42.84 | 58.90 |
| **('Thal', 'Caud', 'Puta', 'Pall', 'Amyg', 'Caud*Puta', 'Caud*Pall', 'Thal*Puta')** | 42.85 | 58.91 |
| **('Thal', 'Caud', 'Puta', 'Pall', 'Hipp', 'Amyg', 'Caud*Puta', 'Caud*Pall')** | 42.85 | 58.91 |
| **('Caud', 'Puta', 'Pall', 'Hipp', 'Amyg', 'Caud*Puta', 'Caud*Pall', 'Pall*Puta')** | 42.88 | 58.94 |
| **('Thal', 'Caud', 'Puta', 'Pall', 'Hipp', 'Amyg', 'Caud*Pall')** | 42.89 | 57.16 |
| **('Thal', 'Caud', 'Puta', 'Pall', 'Hipp', 'Caud*Pall', 'Thal*Puta')** | 42.89 | 57.16 |
| **('Pall', 'Amyg', 'Accu')** | 42.90 | 50.04 |
| **('Thal', 'Caud', 'Puta', 'Pall', 'Accu', 'Caud*Pall', 'Thal*Puta')** | 42.91 | 57.18 |
| **('Caud', 'Puta', 'Pall', 'Hipp', 'Accu', 'Caud*Pall')** | 42.91 | 55.40 |
| **('Caud', 'Puta', 'Pall', 'Amyg', 'Accu', 'Caud*Puta', 'Caud*Pall', 'Pall*Puta', 'Caud*Puta*Pall')** | 42.91 | 60.75 |
| **('Thal', 'Caud', 'Puta', 'Hipp', 'Amyg', 'Accu', 'Caud*Puta', 'Thal*Puta')** | 42.92 | 58.98 |
| **('Thal', 'Caud', 'Puta', 'Pall', 'Amyg', 'Caud*Pall', 'Thal*Puta', 'Caud*Puta*Pall')** | 42.93 | 58.99 |
| **('Thal', 'Caud', 'Puta', 'Pall', 'Hipp', 'Amyg', 'Thal*Puta', 'Caud*Puta*Pall')** | 42.93 | 58.99 |

| | | |
|---|---|---|
| **('Thal', 'Caud', 'Puta', 'Pall', 'Accu', 'Caud*Pall')** | 42.98 | 55.47 |
| **('Thal', 'Caud', 'Puta', 'Pall', 'Accu', 'Pall*Puta', 'Thal*Puta', 'Caud*Puta*Pall')** | 42.98 | 59.04 |
| **('Caud', 'Hipp', 'Amyg')** | 43.00 | 50.13 |
| **('Thal', 'Caud', 'Puta', 'Pall', 'Hipp', 'Caud*Pall')** | 43.00 | 55.49 |
| **('Pall', 'Hipp', 'Amyg')** | 43.01 | 50.15 |
| **('Caud', 'Amyg', 'Accu')** | 43.02 | 50.16 |
| **('Caud', 'Puta', 'Pall', 'Hipp', 'Caud*Pall', 'Caud*Puta*Pall')** | 43.03 | 55.52 |
| **('Thal', 'Caud', 'Puta', 'Pall', 'Hipp', 'Accu', 'Pall*Puta', 'Caud*Puta*Pall')** | 43.03 | 59.09 |
| **('Thal', 'Pall', 'Hipp', 'Amyg')** | 43.04 | 51.96 |
| **('Thal', 'Caud', 'Puta', 'Pall', 'Hipp', 'Amyg', 'Accu')** | 43.05 | 57.32 |
| **('Thal', 'Caud', 'Pall', 'Amyg')** | 43.05 | 51.97 |
| **('Caud', 'Puta', 'Pall', 'Hipp', 'Accu', 'Caud*Puta*Pall')** | 43.05 | 55.54 |
| **('Caud', 'Puta', 'Pall', 'Hipp', 'Amyg', 'Accu', 'Caud*Puta', 'Pall*Puta', 'Caud*Puta*Pall')** | 43.05 | 60.89 |
| **('Thal', 'Caud', 'Puta', 'Pall', 'Hipp', 'Caud*Puta*Pall')** | 43.08 | 55.57 |
| **('Thal', 'Caud', 'Puta', 'Pall', 'Hipp', 'Accu')** | 43.10 | 55.58 |
| **('Thal', 'Caud', 'Puta', 'Pall', 'Caud*Pall', 'Caud*Puta*Pall')** | 43.10 | 55.58 |
| **('Thal', 'Caud', 'Puta', 'Pall', 'Amyg', 'Accu', 'Caud*Puta', 'Pall*Puta', 'Caud*Puta*Pall')** | 43.11 | 60.95 |
| **('Thal', 'Caud', 'Puta', 'Pall', 'Caud*Puta', 'Caud*Pall', 'Thal*Puta', 'Caud*Puta*Pall')** | 43.13 | 59.19 |
| **('Caud', 'Puta', 'Pall', 'Accu', 'Caud*Pall', 'Caud*Puta*Pall')** | 43.14 | 55.63 |
| **('Thal', 'Caud', 'Puta', 'Pall', 'Accu', 'Caud*Puta*Pall')** | 43.14 | 55.63 |
| **('Thal', 'Caud', 'Puta', 'Pall', 'Accu', 'Caud*Puta', 'Thal*Puta', 'Caud*Puta*Pall')** | 43.15 | 59.21 |

| | | |
|---|---|---|
| **('Caud', 'Pall', 'Amyg')** | 43.19 | 50.32 |
| **('Thal', 'Caud', 'Puta', 'Pall', 'Hipp', 'Caud*Puta', 'Thal*Puta', 'Caud*Puta*Pall')** | 43.19 | 59.24 |
| **('Thal', 'Caud', 'Hipp', 'Amyg')** | 43.20 | 52.12 |
| **('Pall', 'Hipp', 'Accu')** | 43.21 | 50.35 |
| **('Caud', 'Puta', 'Pall', 'Hipp', 'Amyg', 'Accu', 'Caud*Pall', 'Pall*Puta', 'Caud*Puta*Pall')** | 43.26 | 61.11 |
| **('Caud', 'Hipp', 'Accu')** | 43.28 | 50.41 |
| **('Caud', 'Puta', 'Pall', 'Hipp', 'Amyg', 'Caud*Puta', 'Caud*Pall', 'Pall*Puta', 'Caud*Puta*Pall')** | 43.28 | 61.12 |
| **('Thal', 'Caud', 'Puta', 'Pall', 'Amyg', 'Caud*Puta', 'Caud*Pall', 'Pall*Puta', 'Caud*Puta*Pall')** | 43.30 | 61.14 |
| **('Thal', 'Caud', 'Puta', 'Pall', 'Amyg', 'Accu', 'Caud*Pall', 'Pall*Puta', 'Caud*Puta*Pall')** | 43.31 | 61.15 |
| **('Thal', 'Caud', 'Puta', 'Pall', 'Hipp', 'Accu', 'Caud*Puta', 'Pall*Puta')** | 43.34 | 59.39 |
| **('Thal', 'Caud', 'Puta', 'Pall', 'Accu', 'Caud*Puta', 'Pall*Puta', 'Thal*Puta')** | 43.39 | 59.44 |
| **('Caud', 'Puta', 'Pall', 'Hipp', 'Accu', 'Caud*Puta', 'Caud*Pall', 'Pall*Puta')** | 43.40 | 59.46 |
| **('Thal', 'Caud', 'Puta', 'Pall', 'Accu', 'Caud*Puta', 'Caud*Pall', 'Pall*Puta')** | 43.40 | 59.46 |
| **('Thal', 'Caud', 'Puta', 'Pall', 'Hipp', 'Caud*Puta', 'Pall*Puta', 'Thal*Puta')** | 43.45 | 59.51 |
| **('Thal', 'Caud', 'Puta', 'Pall', 'Hipp', 'Caud*Puta', 'Caud*Pall', 'Pall*Puta')** | 43.45 | 59.51 |
| **('Thal', 'Caud', 'Puta', 'Pall', 'Hipp', 'Accu', 'Caud*Puta', 'Thal*Puta')** | 43.47 | 59.53 |

| | | |
|---|---|---|
| **('Thal', 'Caud', 'Puta', 'Pall', 'Caud*Puta', 'Caud*Pall', 'Pall*Puta', 'Thal*Puta')** | 43.48 | 59.54 |
| **('Thal', 'Caud', 'Puta', 'Pall', 'Hipp', 'Pall*Puta', 'Thal*Puta', 'Caud*Puta*Pall')** | 43.48 | 59.54 |
| **('Caud', 'Pall', 'Caud*Pall')** | 43.51 | 50.64 |
| **('Caud', 'Pall', 'Hipp')** | 43.52 | 50.65 |
| **('Thal', 'Caud', 'Puta', 'Pall', 'Hipp', 'Amyg', 'Accu', 'Pall*Puta', 'Caud*Puta*Pall')** | 43.53 | 61.37 |
| **('Thal', 'Caud', 'Puta', 'Pall', 'Amyg', 'Accu', 'Pall*Puta', 'Thal*Puta', 'Caud*Puta*Pall')** | 43.55 | 61.39 |
| **('Caud', 'Pall', 'Accu')** | 43.56 | 50.70 |
| **('Thal', 'Caud', 'Puta', 'Pall', 'Hipp', 'Caud*Puta', 'Caud*Pall', 'Thal*Puta')** | 43.60 | 59.66 |
| **('Thal', 'Caud', 'Puta', 'Pall', 'Hipp', 'Amyg', 'Caud*Puta', 'Pall*Puta', 'Caud*Puta*Pall')** | 43.62 | 61.46 |
| **('Thal', 'Caud', 'Puta', 'Pall', 'Hipp', 'Accu', 'Thal*Puta', 'Caud*Puta*Pall')** | 43.63 | 59.69 |
| **('Thal', 'Caud', 'Puta', 'Pall', 'Amyg', 'Caud*Puta', 'Pall*Puta', 'Thal*Puta', 'Caud*Puta*Pall')** | 43.64 | 61.48 |
| **('Thal', 'Caud', 'Puta', 'Pall', 'Accu', 'Caud*Puta', 'Caud*Pall', 'Thal*Puta')** | 43.64 | 59.70 |
| **('Thal', 'Puta', 'Pall', 'Hipp', 'Amyg', 'Accu', 'Pall*Puta', 'Thal*Puta')** | 43.67 | 59.73 |
| **('Thal', 'Caud', 'Puta', 'Pall', 'Amyg', 'Accu', 'Caud*Pall', 'Pall*Puta')** | 43.73 | 59.78 |
| **('Thal', 'Caud', 'Puta', 'Pall', 'Amyg', 'Accu', 'Pall*Puta', 'Thal*Puta')** | 43.73 | 59.79 |
| **('Thal', 'Caud', 'Puta', 'Pall', 'Amyg', 'Accu', 'Caud*Puta', 'Thal*Puta', 'Caud*Puta*Pall')** | 43.73 | 61.58 |

| | | |
|---|---|---|
| **('Thal', 'Caud', 'Puta', 'Pall', 'Hipp', 'Caud*Pall', 'Thal*Puta', 'Caud*Puta*Pall')** | 43.73 | 59.79 |
| **('Thal', 'Caud', 'Puta', 'Pall', 'Amyg', 'Caud*Pall', 'Pall*Puta', 'Thal*Puta', 'Caud*Puta*Pall')** | 43.77 | 61.61 |
| **('Thal', 'Caud', 'Puta', 'Pall', 'Hipp', 'Amyg', 'Accu', 'Pall*Puta')** | 43.78 | 59.84 |
| **('Thal', 'Caud', 'Puta', 'Pall', 'Accu', 'Caud*Pall', 'Thal*Puta', 'Caud*Puta*Pall')** | 43.82 | 59.88 |
| **('Thal', 'Caud', 'Pall', 'Accu')** | 43.83 | 52.76 |
| **('Thal', 'Pall', 'Hipp', 'Accu')** | 43.85 | 52.77 |
| **('Thal', 'Caud', 'Puta', 'Pall', 'Hipp', 'Amyg', 'Accu', 'Thal*Puta')** | 43.85 | 59.91 |
| **('Thal', 'Caud', 'Puta', 'Pall', 'Hipp', 'Amyg', 'Caud*Pall', 'Pall*Puta', 'Caud*Puta*Pall')** | 43.86 | 61.70 |
| **('Thal', 'Caud', 'Hipp', 'Accu')** | 43.87 | 52.79 |
| **('Thal', 'Caud', 'Puta', 'Pall', 'Hipp', 'Accu', 'Caud*Puta', 'Caud*Puta*Pall')** | 43.87 | 59.93 |
| **('Caud', 'Puta', 'Pall', 'Hipp', 'Accu', 'Caud*Puta', 'Caud*Pall', 'Caud*Puta*Pall')** | 43.89 | 59.94 |
| **('Caud', 'Puta', 'Pall', 'Hipp', 'Amyg', 'Accu', 'Caud*Pall', 'Pall*Puta')** | 43.89 | 59.95 |
| **('Thal', 'Caud', 'Puta', 'Pall', 'Amyg', 'Accu', 'Caud*Pall', 'Thal*Puta')** | 43.91 | 59.97 |
| **('Thal', 'Caud', 'Puta', 'Pall', 'Hipp', 'Caud*Puta', 'Caud*Pall', 'Caud*Puta*Pall')** | 43.94 | 60.00 |
| **('Caud', 'Puta', 'Pall', 'Hipp', 'Accu', 'Caud*Puta', 'Caud*Pall', 'Pall*Puta', 'Caud*Puta*Pall')** | 43.96 | 61.80 |
| **('Thal', 'Caud', 'Puta', 'Pall', 'Amyg', 'Accu', 'Caud*Pall', 'Caud*Puta*Pall')** | 43.97 | 60.03 |
| **('Thal', 'Caud', 'Puta', 'Pall', 'Hipp', 'Amyg', 'Pall*Puta', 'Thal*Puta')** | 43.98 | 60.03 |

| | | |
|---|---|---|
| **('Thal', 'Caud', 'Puta', 'Pall', 'Accu', 'Caud*Puta', 'Caud*Pall', 'Caud*Puta*Pall')** | 43.98 | 60.04 |
| **('Thal', 'Caud', 'Puta', 'Pall', 'Hipp', 'Accu', 'Caud*Puta', 'Caud*Pall')** | 43.98 | 60.04 |
| **('Thal', 'Caud', 'Puta', 'Pall', 'Accu', 'Caud*Puta', 'Caud*Pall', 'Pall*Puta', 'Caud*Puta*Pall')** | 44.00 | 61.84 |
| **('Thal', 'Caud', 'Puta', 'Pall', 'Hipp', 'Amyg', 'Caud*Puta', 'Thal*Puta', 'Caud*Puta*Pall')** | 44.01 | 61.85 |
| **('Caud', 'Puta', 'Pall', 'Hipp', 'Amyg', 'Accu', 'Caud*Pall', 'Caud*Puta*Pall')** | 44.01 | 60.07 |
| **('Thal', 'Caud', 'Puta', 'Pall', 'Amyg', 'Caud*Puta', 'Caud*Pall', 'Thal*Puta', 'Caud*Puta*Pall')** | 44.01 | 61.86 |
| **('Thal', 'Caud', 'Puta', 'Pall', 'Amyg', 'Caud*Pall', 'Pall*Puta', 'Thal*Puta')** | 44.02 | 60.07 |
| **('Thal', 'Caud', 'Pall', 'Caud*Pall')** | 44.02 | 52.94 |
| **('Thal', 'Caud', 'Puta', 'Pall', 'Hipp', 'Amyg', 'Caud*Pall', 'Pall*Puta')** | 44.06 | 60.12 |
| **('Thal', 'Caud', 'Puta', 'Pall', 'Hipp', 'Amyg', 'Accu', 'Caud*Puta', 'Caud*Puta*Pall')** | 44.08 | 61.92 |
| **('Caud', 'Puta', 'Pall', 'Hipp', 'Amyg', 'Accu', 'Caud*Puta', 'Caud*Pall', 'Caud*Puta*Pall')** | 44.10 | 61.95 |
| **('Thal', 'Caud', 'Puta', 'Pall', 'Amyg', 'Accu', 'Caud*Puta', 'Caud*Pall', 'Caud*Puta*Pall')** | 44.11 | 61.95 |
| **('Thal', 'Caud', 'Pall', 'Hipp')** | 44.14 | 53.06 |
| **('Thal', 'Caud', 'Puta', 'Pall', 'Caud*Puta', 'Caud*Pall', 'Pall*Puta', 'Thal*Puta', 'Caud*Puta*Pall')** | 44.14 | 61.98 |
| **('Thal', 'Caud', 'Puta', 'Pall', 'Hipp', 'Amyg', 'Caud*Pall', 'Thal*Puta')** | 44.15 | 60.21 |

| | | |
|---|---|---|
| **('Thal', 'Caud', 'Puta', 'Pall', 'Hipp', 'Amyg', 'Accu', 'Caud*Puta*Pall')** | 44.16 | 60.21 |
| **('Thal', 'Caud', 'Puta', 'Pall', 'Hipp', 'Caud*Puta', 'Caud*Pall', 'Pall*Puta', 'Caud*Puta*Pall')** | 44.18 | 62.02 |
| **('Thal', 'Caud', 'Puta', 'Pall', 'Hipp', 'Amyg', 'Accu', 'Thal*Puta', 'Caud*Puta*Pall')** | 44.22 | 62.07 |
| **('Thal', 'Caud', 'Puta', 'Pall', 'Accu', 'Caud*Pall', 'Pall*Puta', 'Thal*Puta', 'Caud*Puta*Pall')** | 44.24 | 62.08 |
| **('Thal', 'Caud', 'Puta', 'Pall', 'Hipp', 'Amyg', 'Pall*Puta', 'Thal*Puta', 'Caud*Puta*Pall')** | 44.31 | 62.15 |
| **('Thal', 'Caud', 'Puta', 'Pall', 'Hipp', 'Amyg', 'Caud*Puta', 'Caud*Pall', 'Caud*Puta*Pall')** | 44.31 | 62.16 |
| **('Thal', 'Caud', 'Puta', 'Pall', 'Hipp', 'Accu', 'Pall*Puta', 'Thal*Puta')** | 44.32 | 60.38 |
| **('Thal', 'Caud', 'Puta', 'Pall', 'Hipp', 'Amyg', 'Caud*Pall', 'Caud*Puta*Pall')** | 44.39 | 60.44 |
| **('Thal', 'Caud', 'Puta', 'Pall', 'Hipp', 'Accu', 'Caud*Puta', 'Pall*Puta', 'Caud*Puta*Pall')** | 44.39 | 62.23 |
| **('Thal', 'Caud', 'Puta', 'Pall', 'Hipp', 'Accu', 'Caud*Pall', 'Pall*Puta', 'Caud*Puta*Pall')** | 44.43 | 62.27 |
| **('Thal', 'Caud', 'Puta', 'Pall', 'Accu', 'Caud*Puta', 'Pall*Puta', 'Thal*Puta', 'Caud*Puta*Pall')** | 44.45 | 62.29 |
| **('Thal', 'Caud', 'Puta', 'Pall', 'Amyg', 'Accu', 'Caud*Pall', 'Thal*Puta', 'Caud*Puta*Pall')** | 44.45 | 62.29 |
| **('Thal', 'Caud', 'Puta', 'Pall', 'Accu', 'Caud*Pall', 'Pall*Puta', 'Thal*Puta')** | 44.46 | 60.52 |
| **('Thal', 'Caud', 'Puta', 'Pall', 'Hipp', 'Amyg', 'Accu', 'Caud*Pall')** | 44.48 | 60.53 |

| | | |
|---|---|---|
| **('Thal', 'Caud', 'Puta', 'Pall', 'Hipp', 'Caud*Pall', 'Pall*Puta', 'Thal*Puta', 'Caud*Puta*Pall')** | 44.49 | 62.33 |
| **('Thal', 'Caud', 'Puta', 'Pall', 'Amyg', 'Accu', 'Caud*Puta', 'Caud*Pall', 'Pall*Puta')** | 44.55 | 62.40 |
| **('Thal', 'Caud', 'Pall', 'Amyg', 'Accu')** | 44.57 | 55.27 |
| **('Thal', 'Pall', 'Hipp', 'Amyg', 'Accu')** | 44.57 | 55.27 |
| **('Thal', 'Caud', 'Puta', 'Pall', 'Hipp', 'Accu', 'Caud*Pall', 'Pall*Puta')** | 44.58 | 60.63 |
| **('Thal', 'Caud', 'Puta', 'Pall', 'Hipp', 'Amyg', 'Accu', 'Caud*Puta', 'Pall*Puta')** | 44.58 | 62.42 |
| **('Thal', 'Caud', 'Puta', 'Pall', 'Hipp', 'Caud*Pall', 'Pall*Puta', 'Thal*Puta')** | 44.58 | 60.64 |
| **('Caud', 'Hipp', 'Amyg', 'Accu')** | 44.60 | 53.52 |
| **('Pall', 'Hipp', 'Amyg', 'Accu')** | 44.60 | 53.52 |
| **('Thal', 'Caud', 'Puta', 'Pall', 'Amyg', 'Accu', 'Caud*Puta', 'Pall*Puta', 'Thal*Puta')** | 44.62 | 62.46 |
| **('Thal', 'Caud', 'Puta', 'Pall', 'Hipp', 'Accu', 'Caud*Pall', 'Thal*Puta')** | 44.62 | 60.68 |
| **('Thal', 'Caud', 'Puta', 'Pall', 'Hipp', 'Amyg', 'Accu', 'Caud*Puta', 'Caud*Pall')** | 44.64 | 62.49 |
| **('Thal', 'Caud', 'Puta', 'Pall', 'Amyg', 'Accu', 'Caud*Puta', 'Caud*Pall', 'Thal*Puta')** | 44.66 | 62.50 |
| **('Thal', 'Caud', 'Puta', 'Pall', 'Hipp', 'Amyg', 'Accu', 'Caud*Puta', 'Thal*Puta')** | 44.68 | 62.52 |
| **('Caud', 'Puta', 'Pall', 'Hipp', 'Amyg', 'Accu', 'Caud*Puta', 'Caud*Pall', 'Pall*Puta')** | 44.68 | 62.52 |
| **('Thal', 'Caud', 'Hipp', 'Amyg', 'Accu')** | 44.71 | 55.41 |
| **('Thal', 'Caud', 'Puta', 'Pall', 'Hipp', 'Amyg', 'Caud*Puta', 'Caud*Pall', 'Pall*Puta')** | 44.73 | 62.57 |

| | | |
|---|---|---|
| **('Thal', 'Caud', 'Puta', 'Pall', 'Hipp', 'Amyg', 'Caud*Pall', 'Thal*Puta', 'Caud*Puta*Pall')** | 44.74 | 62.58 |
| **('Thal', 'Caud', 'Puta', 'Pall', 'Hipp', 'Amyg', 'Caud*Puta', 'Pall*Puta', 'Thal*Puta')** | 44.74 | 62.58 |
| **('Thal', 'Caud', 'Puta', 'Pall', 'Amyg', 'Caud*Puta', 'Caud*Pall', 'Pall*Puta', 'Thal*Puta')** | 44.75 | 62.59 |
| **('Thal', 'Caud', 'Puta', 'Pall', 'Hipp', 'Caud*Puta', 'Pall*Puta', 'Thal*Puta', 'Caud*Puta*Pall')** | 44.75 | 62.59 |
| **('Thal', 'Caud', 'Puta', 'Pall', 'Hipp', 'Accu', 'Pall*Puta', 'Thal*Puta', 'Caud*Puta*Pall')** | 44.79 | 62.63 |
| **('Caud', 'Pall', 'Amyg', 'Caud*Pall')** | 44.80 | 53.72 |
| **('Thal', 'Caud', 'Puta', 'Pall', 'Hipp', 'Amyg', 'Caud*Puta', 'Caud*Pall', 'Thal*Puta')** | 44.81 | 62.65 |
| **('Thal', 'Caud', 'Puta', 'Pall', 'Hipp', 'Accu', 'Caud*Pall')** | 44.82 | 59.09 |
| **('Thal', 'Caud', 'Pall', 'Amyg', 'Caud*Pall')** | 44.84 | 55.55 |
| **('Caud', 'Puta', 'Pall', 'Hipp', 'Accu', 'Caud*Pall', 'Caud*Puta*Pall')** | 44.84 | 59.12 |
| **('Caud', 'Puta', 'Pall', 'Hipp', 'Amyg', 'Accu', 'Caud*Puta', 'Caud*Pall', 'Pall*Puta', 'Caud*Puta*Pall')** | 44.86 | 64.48 |
| **('Thal', 'Caud', 'Puta', 'Pall', 'Hipp', 'Accu', 'Caud*Puta', 'Thal*Puta', 'Caud*Puta*Pall')** | 44.88 | 62.72 |
| **('Caud', 'Pall', 'Amyg', 'Accu')** | 44.88 | 53.80 |
| **('Thal', 'Caud', 'Puta', 'Pall', 'Hipp', 'Caud*Puta', 'Caud*Pall', 'Thal*Puta', 'Caud*Puta*Pall')** | 44.91 | 62.75 |
| **('Thal', 'Caud', 'Puta', 'Pall', 'Amyg', 'Accu', 'Caud*Puta', 'Caud*Pall', 'Pall*Puta', 'Caud*Puta*Pall')** | 44.91 | 64.53 |

| | | |
|---|---|---|
| **('Thal', 'Caud', 'Puta', 'Pall', 'Accu', 'Caud*Pall', 'Caud*Puta*Pall')** | 44.95 | 59.23 |
| **('Caud', 'Pall', 'Hipp', 'Amyg')** | 44.96 | 53.88 |
| **('Thal', 'Caud', 'Puta', 'Pall', 'Hipp', 'Caud*Pall', 'Caud*Puta*Pall')** | 44.97 | 59.25 |
| **('Thal', 'Caud', 'Puta', 'Pall', 'Hipp', 'Accu', 'Caud*Puta*Pall')** | 44.98 | 59.26 |
| **('Thal', 'Caud', 'Puta', 'Pall', 'Accu', 'Caud*Puta', 'Caud*Pall', 'Thal*Puta', 'Caud*Puta*Pall')** | 45.00 | 62.84 |
| **('Thal', 'Caud', 'Pall', 'Hipp', 'Amyg')** | 45.04 | 55.75 |
| **('Thal', 'Caud', 'Puta', 'Pall', 'Hipp', 'Amyg', 'Accu', 'Caud*Puta', 'Pall*Puta', 'Caud*Puta*Pall')** | 45.05 | 64.68 |
| **('Caud', 'Pall', 'Accu', 'Caud*Pall')** | 45.05 | 53.97 |
| **('Thal', 'Caud', 'Puta', 'Pall', 'Amyg', 'Accu', 'Caud*Puta', 'Pall*Puta', 'Thal*Puta', 'Caud*Puta*Pall')** | 45.11 | 64.73 |
| **('Caud', 'Pall', 'Hipp', 'Accu')** | 45.21 | 54.13 |
| **('Thal', 'Caud', 'Puta', 'Pall', 'Amyg', 'Accu', 'Caud*Pall', 'Pall*Puta', 'Thal*Puta', 'Caud*Puta*Pall')** | 45.22 | 64.85 |
| **('Thal', 'Caud', 'Puta', 'Pall', 'Hipp', 'Amyg', 'Accu', 'Caud*Pall', 'Pall*Puta', 'Caud*Puta*Pall')** | 45.26 | 64.89 |
| **('Caud', 'Pall', 'Hipp', 'Caud*Pall')** | 45.28 | 54.20 |
| **('Thal', 'Caud', 'Puta', 'Pall', 'Hipp', 'Amyg', 'Caud*Puta', 'Caud*Pall', 'Pall*Puta', 'Caud*Puta*Pall')** | 45.28 | 64.91 |
| **('Thal', 'Caud', 'Puta', 'Pall', 'Amyg', 'Caud*Puta', 'Caud*Pall', 'Pall*Puta', 'Thal*Puta', 'Caud*Puta*Pall')** | 45.29 | 64.92 |
| **('Thal', 'Caud', 'Puta', 'Pall', 'Hipp', 'Accu', 'Caud*Puta', 'Pall*Puta', 'Thal*Puta')** | 45.30 | 63.14 |

| | | |
|---|---|---|
| **('Thal', 'Caud', 'Puta', 'Pall', 'Hipp', 'Accu', 'Caud*Puta', 'Caud*Pall', 'Pall*Puta')** | 45.33 | 63.18 |
| **('Thal', 'Caud', 'Puta', 'Pall', 'Hipp', 'Accu', 'Caud*Pall', 'Thal*Puta', 'Caud*Puta*Pall')** | 45.36 | 63.20 |
| **('Thal', 'Caud', 'Puta', 'Pall', 'Accu', 'Caud*Puta', 'Caud*Pall', 'Pall*Puta', 'Thal*Puta')** | 45.38 | 63.22 |
| **('Thal', 'Caud', 'Puta', 'Pall', 'Hipp', 'Caud*Puta', 'Caud*Pall', 'Pall*Puta', 'Thal*Puta')** | 45.42 | 63.26 |
| **('Thal', 'Caud', 'Puta', 'Pall', 'Hipp', 'Amyg', 'Accu', 'Pall*Puta', 'Thal*Puta', 'Caud*Puta*Pall')** | 45.45 | 65.08 |
| **('Thal', 'Caud', 'Puta', 'Pall', 'Hipp', 'Accu', 'Caud*Puta', 'Caud*Pall', 'Thal*Puta')** | 45.47 | 63.32 |
| **('Thal', 'Caud', 'Pall', 'Accu', 'Caud*Pall')** | 45.52 | 56.23 |
| **('Thal', 'Caud', 'Puta', 'Pall', 'Hipp', 'Amyg', 'Accu', 'Caud*Puta', 'Thal*Puta', 'Caud*Puta*Pall')** | 45.57 | 65.20 |
| **('Thal', 'Caud', 'Puta', 'Pall', 'Hipp', 'Amyg', 'Caud*Puta', 'Pall*Puta', 'Thal*Puta', 'Caud*Puta*Pall')** | 45.62 | 65.25 |
| **('Thal', 'Caud', 'Puta', 'Pall', 'Hipp', 'Amyg', 'Accu', 'Pall*Puta', 'Thal*Puta')** | 45.65 | 63.49 |
| **('Thal', 'Caud', 'Puta', 'Pall', 'Amyg', 'Accu', 'Caud*Pall', 'Pall*Puta', 'Thal*Puta')** | 45.67 | 63.51 |
| **('Thal', 'Caud', 'Puta', 'Pall', 'Hipp', 'Amyg', 'Accu', 'Caud*Pall', 'Pall*Puta')** | 45.68 | 63.52 |
| **('Thal', 'Caud', 'Puta', 'Pall', 'Amyg', 'Accu', 'Caud*Puta', 'Caud*Pall', 'Thal*Puta', 'Caud*Puta*Pall')** | 45.72 | 65.34 |

| | | |
|---|---|---|
| **('Thal', 'Caud', 'Puta', 'Pall', 'Hipp', 'Amyg', 'Caud*Pall', 'Pall*Puta', 'Thal*Puta', 'Caud*Puta*Pall')** | 45.73 | 65.35 |
| **('Thal', 'Caud', 'Puta', 'Pall', 'Hipp', 'Amyg', 'Accu', 'Caud*Pall', 'Thal*Puta')** | 45.75 | 63.59 |
| **('Thal', 'Caud', 'Pall', 'Hipp', 'Accu')** | 45.82 | 56.52 |
| **('Thal', 'Caud', 'Puta', 'Pall', 'Hipp', 'Accu', 'Caud*Puta', 'Caud*Pall', 'Caud*Puta*Pall')** | 45.87 | 63.71 |
| **('Thal', 'Caud', 'Puta', 'Pall', 'Hipp', 'Amyg', 'Accu', 'Caud*Pall', 'Caud*Puta*Pall')** | 45.88 | 63.72 |
| **('Thal', 'Caud', 'Puta', 'Pall', 'Accu', 'Caud*Puta', 'Caud*Pall', 'Pall*Puta', 'Thal*Puta', 'Caud*Puta*Pall')** | 45.91 | 65.53 |
| **('Thal', 'Caud', 'Puta', 'Pall', 'Hipp', 'Amyg', 'Caud*Puta', 'Caud*Pall', 'Thal*Puta', 'Caud*Puta*Pall')** | 45.91 | 65.54 |
| **('Thal', 'Caud', 'Puta', 'Pall', 'Hipp', 'Accu', 'Caud*Puta', 'Caud*Pall', 'Pall*Puta', 'Caud*Puta*Pall')** | 45.93 | 65.56 |
| **('Thal', 'Caud', 'Puta', 'Pall', 'Hipp', 'Amyg', 'Caud*Pall', 'Pall*Puta', 'Thal*Puta')** | 45.97 | 63.81 |
| **('Thal', 'Caud', 'Pall', 'Amyg', 'Accu', 'Caud*Pall')** | 46.02 | 58.51 |
| **('Thal', 'Caud', 'Pall', 'Hipp', 'Caud*Pall')** | 46.02 | 56.72 |
| **('Thal', 'Caud', 'Puta', 'Pall', 'Hipp', 'Amyg', 'Accu', 'Caud*Puta', 'Caud*Pall', 'Caud*Puta*Pall')** | 46.04 | 65.66 |
| **('Thal', 'Caud', 'Puta', 'Pall', 'Hipp', 'Caud*Puta', 'Caud*Pall', 'Pall*Puta', 'Thal*Puta', 'Caud*Puta*Pall')** | 46.06 | 65.69 |
| **('Thal', 'Caud', 'Puta', 'Pall', 'Hipp', 'Accu', 'Caud*Pall', 'Pall*Puta', 'Thal*Puta', 'Caud*Puta*Pall')** | 46.07 | 65.70 |

| | | |
|---|---|---|
| ('Caud', 'Pall', 'Amyg', 'Accu', 'Caud*Pall') | 46.12 | 56.82 |
| ('Thal', 'Caud', 'Puta', 'Pall', 'Hipp', 'Amyg', 'Accu', 'Caud*Pall', 'Thal*Puta', 'Caud*Puta*Pall') | 46.16 | 65.79 |
| ('Thal', 'Caud', 'Puta', 'Pall', 'Hipp', 'Accu', 'Caud*Pall', 'Pall*Puta', 'Thal*Puta') | 46.32 | 64.16 |
| ('Thal', 'Caud', 'Puta', 'Pall', 'Hipp', 'Accu', 'Caud*Puta', 'Pall*Puta', 'Thal*Puta', 'Caud*Puta*Pall') | 46.32 | 65.95 |
| ('Thal', 'Caud', 'Puta', 'Pall', 'Hipp', 'Amyg', 'Accu', 'Caud*Puta', 'Caud*Pall', 'Pall*Puta') | 46.51 | 66.13 |
| ('Caud', 'Pall', 'Hipp', 'Amyg', 'Accu') | 46.55 | 57.25 |
| ('Thal', 'Caud', 'Puta', 'Pall', 'Amyg', 'Accu', 'Caud*Puta', 'Caud*Pall', 'Pall*Puta', 'Thal*Puta') | 46.55 | 66.18 |
| ('Thal', 'Caud', 'Pall', 'Hipp', 'Amyg', 'Accu') | 46.57 | 59.06 |
| ('Thal', 'Caud', 'Puta', 'Pall', 'Hipp', 'Amyg', 'Accu', 'Caud*Puta', 'Pall*Puta', 'Thal*Puta') | 46.57 | 66.20 |
| ('Thal', 'Caud', 'Puta', 'Pall', 'Hipp', 'Amyg', 'Accu', 'Caud*Puta', 'Caud*Pall', 'Thal*Puta') | 46.59 | 66.21 |
| ('Caud', 'Pall', 'Hipp', 'Amyg', 'Caud*Pall') | 46.60 | 57.31 |
| ('Thal', 'Caud', 'Puta', 'Pall', 'Hipp', 'Accu', 'Caud*Puta', 'Caud*Pall', 'Thal*Puta', 'Caud*Puta*Pall') | 46.69 | 66.32 |
| ('Caud', 'Pall', 'Hipp', 'Accu', 'Caud*Pall') | 46.70 | 57.40 |
| ('Thal', 'Caud', 'Puta', 'Pall', 'Hipp', 'Amyg', 'Caud*Puta', 'Caud*Pall', 'Pall*Puta', 'Thal*Puta') | 46.73 | 66.35 |
| ('Thal', 'Caud', 'Puta', 'Pall', 'Hipp', 'Accu', 'Caud*Pall', 'Caud*Puta*Pall') | 46.78 | 62.84 |

| | | |
|---|---|---|
| **('Thal', 'Caud', 'Pall', 'Hipp', 'Amyg', 'Caud*Pall')** | 46.83 | 59.32 |
| **('Thal', 'Caud', 'Puta', 'Pall', 'Hipp', 'Amyg', 'Accu', 'Caud*Puta', 'Caud*Pall', 'Pall*Puta', 'Caud*Puta*Pall')** | 46.86 | 68.27 |
| **('Thal', 'Caud', 'Puta', 'Pall', 'Amyg', 'Accu', 'Caud*Puta', 'Caud*Pall', 'Pall*Puta', 'Thal*Puta', 'Caud*Puta*Pall')** | 46.89 | 68.30 |
| **('Thal', 'Caud', 'Puta', 'Pall', 'Hipp', 'Amyg', 'Accu', 'Caud*Puta', 'Pall*Puta', 'Thal*Puta', 'Caud*Puta*Pall')** | 47.04 | 68.45 |
| **('Thal', 'Caud', 'Puta', 'Pall', 'Hipp', 'Amyg', 'Accu', 'Caud*Pall', 'Pall*Puta', 'Thal*Puta', 'Caud*Puta*Pall')** | 47.12 | 68.53 |
| **('Thal', 'Caud', 'Puta', 'Pall', 'Hipp', 'Amyg', 'Caud*Puta', 'Caud*Pall', 'Pall*Puta', 'Thal*Puta', 'Caud*Puta*Pall')** | 47.26 | 68.67 |
| **('Thal', 'Caud', 'Puta', 'Pall', 'Hipp', 'Accu', 'Caud*Puta', 'Caud*Pall', 'Pall*Puta', 'Thal*Puta')** | 47.29 | 66.92 |
| **('Thal', 'Caud', 'Pall', 'Hipp', 'Accu', 'Caud*Pall')** | 47.50 | 59.99 |
| **('Thal', 'Caud', 'Puta', 'Pall', 'Hipp', 'Amyg', 'Accu', 'Caud*Puta', 'Caud*Pall', 'Thal*Puta', 'Caud*Puta*Pall')** | 47.54 | 68.95 |
| **('Thal', 'Caud', 'Puta', 'Pall', 'Hipp', 'Amyg', 'Accu', 'Caud*Pall', 'Pall*Puta', 'Thal*Puta')** | 47.58 | 67.21 |
| **('Thal', 'Caud', 'Puta', 'Pall', 'Hipp', 'Accu', 'Caud*Puta', 'Caud*Pall', 'Pall*Puta', 'Thal*Puta', 'Caud*Puta*Pall')** | 47.78 | 69.19 |
| **('Caud', 'Pall', 'Hipp', 'Amyg', 'Accu', 'Caud*Pall')** | 47.79 | 60.28 |
| **('Thal', 'Caud', 'Pall', 'Hipp', 'Amyg', 'Accu', 'Caud*Pall')** | 48.01 | 62.29 |

| | | |
|---|---|---|
| **('Thal', 'Caud', 'Puta', 'Pall', 'Hipp', 'Amyg', 'Accu', 'Caud*Puta', 'Caud*Pall', 'Pall*Puta', 'Thal*Puta')** | 48.51 | 69.92 |
| **('Thal', 'Caud', 'Puta', 'Pall', 'Hipp', 'Amyg', 'Accu', 'Caud*Puta', 'Caud*Pall', 'Pall*Puta', 'Thal*Puta', 'Caud*Puta*Pall')** | 48.82 | 72.02 |

Table 1. All the possible combinations of subcortical structures to predict the landmark point of subjective equality (PSE) and their corresponding AIC and BIC. As it is shown on the first row, the model containing Putamen has the lowest AIC and BIC, making it the winning model. Thal: thalamus, Caud: Caudate, Puta: Putamen, Pall: Globus Pallidus, Hipp: Hippocampus, Amyg: Amygdala, Accu: Nucleus accumbense.

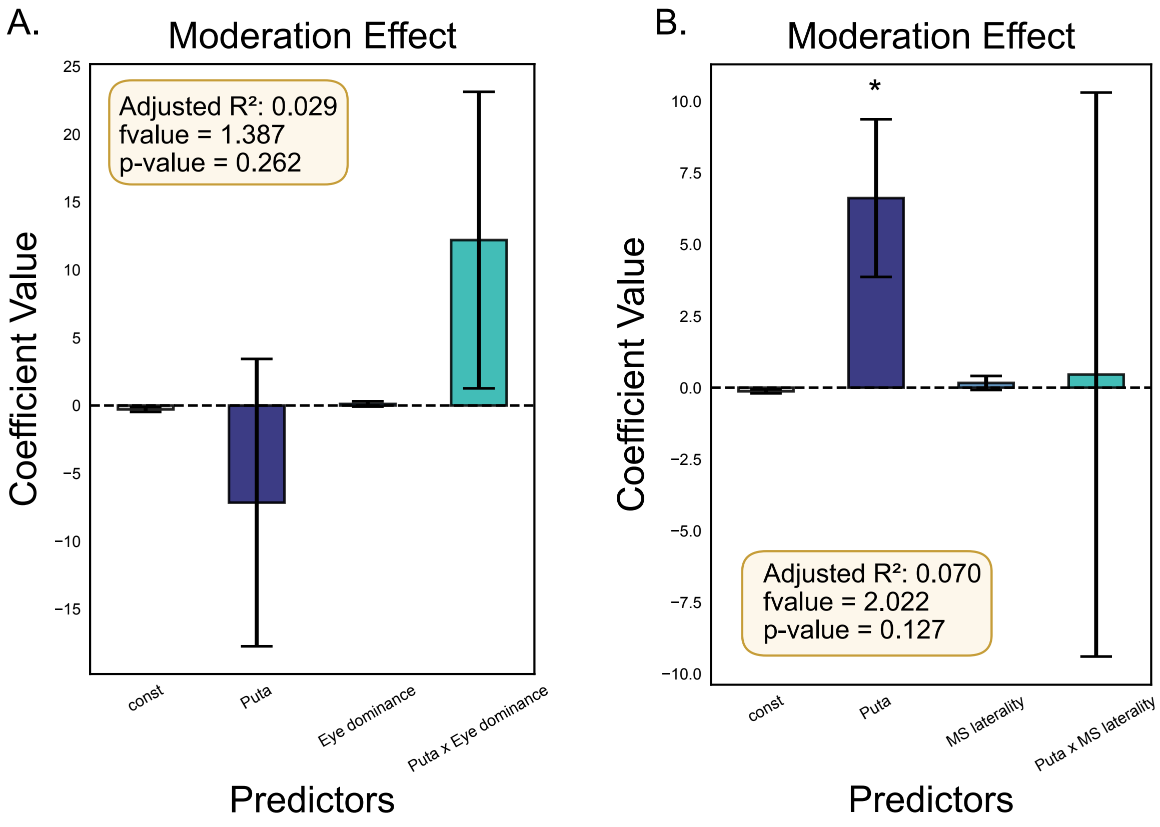


Figure 1. Moderation effect of eye dominance and microsaccade laterality on the relationship between putamen asymmetry and landmark PSE. A) The regression model which included eye dominance, putamen asymmetry, and their interaction, did not show a significant association with landmark PSE scores, suggesting that the strength of the putamen-attention bias relationship does not vary as a function of eye dominance. Adjusted $R^2 = 0.029$, model *p*

= 0.262. B) The regression model which included microsaccade laterality, putamen asymmetry, and their interaction, did not show a significant association with landmark PSE scores, suggesting that the strength of the putamen-attention bias relationship does not vary as a function of microsaccade laterality. Adjusted $R^2 = 0.070$, model $p = 0.127$. Puta = Putamen, MS = Microsaccade; asterisks (*) denotes $p < 0.05$.

A.

| Coefficient | Coefficient Value | Standard Error | 95% Confidence Interval | p-value |
|---|---|---|---|---|
| Constant | -0.291 | 0.181 | [-0.645, 0.063] | 0.116 |
| Putamen | -7.16 | 10.596 | [-27.928, 13.607] | 0.504 |
| Eye dominance | 0.113 | 0.196 | [-0.271, 0.496] | 0.568 |
| Putamen- Eye dominance interaction | 12.197 | 10.934 | [-9.234, 33.627] | 0.272 |

B.

| Coefficient | Coefficient Value | Standard Error | 95% Confidence Interval | p-value |
|---|---|---|---|---|
| Constant | -0.13 | 0.071 | [-0.27, 0.01] | 0.077 |
| Putamen | 6.622 | 2.753 | [1.225, 12.019] | 0.021 |
| MS laterality | 0.164 | 0.245 | [-0.316, 0.644] | 0.508 |
| Putamen- MS laterality interaction | 0.458 | 9.854 | [-18.856, 19.771] | 0.963 |

Table 2. coefficient estimates of moderation analysis of eye dominance (A) and microsaccade laterality (B). MS = Microsaccade
